# Longitudinal single-cell chemical imaging of engineered strains reveals heterogeneity in fatty acid production

**DOI:** 10.1101/2021.07.26.453865

**Authors:** Nathan Tague, Haonan Lin, Jean-Baptiste Lugagne, Owen M. O’Connor, Deeya Burman, Wilson W. Wong, Ji-Xin Cheng, Mary J. Dunlop

## Abstract

Understanding metabolic heterogeneity is critical for optimizing microbial production of valuable chemicals, but requires tools that can quantify metabolites at the single-cell level over time. Here, we develop longitudinal hyperspectral stimulated Raman scattering (SRS) chemical imaging to directly visualize free fatty acids in engineered *Escherichia coli* over many cell cycles. We also develop compositional analysis to determine the chain length and unsaturation of the fatty acids in living cells. Our method reveals substantial heterogeneity in fatty acid production among and within colonies that emerges over the course of many generations. Interestingly, the strains display distinct types of production heterogeneity in an enzyme-dependent manner. By pairing time-lapse and SRS imaging, we examine the relationship between growth and production at the single-cell level. Single-cell quantification does not show a significant growth-production tradeoff in a strain that exhibits high production heterogeneity. Our results demonstrate that cell-to-cell production heterogeneity is pervasive and provide a means to link single-cell and population-level production.

## Introduction

Microbial production of chemicals has the potential to provide a sustainable source of products ranging from fuels to specialty materials (1–4). A major difficulty holding back the replacement of industrial chemicals with bio-based alternatives is that bioproduction often falls short in terms of conversion metrics that dictate economic feasibility, such as titer, rate, and yield. Over the past two decades, researchers have made great strides in identifying metabolic pathways capable of producing a diverse array of useful chemicals (5). However, the reality is that extensive engineering and optimization are required for any given chemical to compete as an alternative to those sourced from petroleum.

Producing chemicals in cells offers many advantages, but presents unique industrial challenges. For example, cell-to-cell variation and genetic mutations can result in production heterogeneity during fermentation that limits overall process efficiency. Single-cell variation can stem from a variety of causes, such as stochasticity in the underlying biological processes (6, 7), variations in media environments within cultures (8), or selection pressures against high producing cells causing mutational escape variants (9, 10). However, the frequency and impact of production variation and how it changes over time are largely unknown. Methods that enable quantification of heterogeneity and its emergence are a prerequisite to understanding the root cause and implementing designs that mitigate its effect on overall efficiency.

Here, we focus on fatty acid synthesis, which is an attractive pathway for metabolic engineering because it offers a biological means to synthesize linear hydrocarbons. Fatty acids and their derivatives are high demand chemicals that can be used as fuels, commodities, and specialty chemicals. Numerous studies have aimed at increasing the efficiency of fatty acid synthesis pathways as well as controlling the species of fatty acid produced (11–14). Termination enzymes that interface with this pathway can be used to produce a wide variety of high-value fatty acid derivatives such as alkanes, olefins, and alcohols (15).

Current methods to measure production strain performance include mass spectrometry, fluorescent biosensors, and dyes. Mass spectrometry-based techniques provide exquisite chemical specificity but are limited in their ability to quantify single cells, which means they can overlook valuable information about population heterogeneity that is key to predicting population stability during scale-up (16–18). Further, because the measurement process is destructive, it is not possible to follow production changes within the same cells over time. Biosensor-based fluorescent assays, in contrast, can capture dynamic, single-cell information. These systems are amenable to high throughput screens and are non-destructive (19). However, well-characterized biosensors are scarce in comparison to the number of chemicals metabolic engineers can produce. Additionally, significant optimization is often necessary to fine tune the concentration responsive range of a biosensor (20–22). In the case of fatty acid production, lipophilic dyes such as Nile red have been used to measure production (23), however these stains lack lipid specificity. Further, both biosensor and dye-based measurements are indirect readouts of chemical production.

Given the drawbacks of current screening methods, we sought to develop an alternative approach that can capture production and composition information in single cells over time. Stimulated Raman scattering (SRS) is an ideal candidate, as it is a non-destructive, label-free vibrational spectroscopic imaging method that directly detects chemical compounds based on intrinsic molecular vibrations (24, 25). The ability of SRS to probe metabolic activities in live cells has been demonstrated on microalgae (26) and mammalian cells (27) for short periods of time. Although SRS imaging of industrially relevant microbes such as *E. coli* has been reported (28, 29), its use has been limited to conditions where cells were either fixed or where only a single timepoint was required. Performing longitudinal SRS for compositional chemical imaging on live microbes remains challenging. This is mainly attributed to their small size (e.g. *E. coli* are 1-2 μm in length), which shortens the axial signal integration length, and thus yields weaker SRS signals compared to larger cells. In the context of metabolic engineering, where compositional information on products is critical, one needs to perform hyperspectral SRS to generate pixel-wise Raman spectra for molecular fingerprinting. However, due to significant spectral overlaps between metabolites, especially in the carbon-hydrogen (C-H) region, existing hyperspectral SRS image processing methods only provide unsaturation levels of fatty acids (30). They also fail to deliver information on chain length, which is equally important for free fatty acid synthesis.

Here, we introduce a longitudinal hyperspectral SRS method to study metabolically engineered *E. coli*, monitoring free fatty acid production and composition in live cells. We perform SRS in the C-H region which maximizes SRS signals. To overcome spectral cross-talk in the region, we develop a hyperspectral image analysis technique that generates chain length and unsaturation level predictions, allowing for chemical readouts that are analogous to GC-MS. First, we demonstrate that we can clearly distinguish fatty acid production strains from wild type *E. coli* by deconstructing images into maps of their chemical components. With the ability to measure production at the single-cell level, we examine heterogeneity in fatty acid production strains and observe both colony-level heterogeneity and substantial cell-to-cell differences in production. We optimize imaging parameters to enable longitudinal hyperspectral SRS imaging to capture fatty acid production over time in growing cells. Next, we use longitudinal measurements to demonstrate dynamic differences in fatty acid production and composition within the same strain. To the best of our knowledge, this is the first demonstration of longitudinal hyperspectral SRS imaging of live cells over many cell cycles. Lastly, we pair SRS microscopy with time-lapse phase contrast microscopy and automated segmentation analysis to examine relationships between production and growth.

Overall, our study presents two important advances of SRS microscopy, namely fatty acid chain length extraction and longitudinal imaging of proliferating cells. Upon these advances, we report discoveries of metabolic heterogeneity among different cells in a colony and temporal heterogeneity throughout colony formation.

## Results

### Hyperspectral SRS imaging of fatty acid production strains

Spectral signals from Raman scattering correspond to vibrational energies of covalent bonds. This allows for direct imaging of chemicals without the need for labels such as fluorescent reporters or dyes. Here, we deploy hyperspectral SRS (31–33) to obtain chemical maps of protein and fatty acids. To achieve this, we chirp two broadband femtosecond laser beams (pump and Stokes) using high-dispersion glass rods, producing linear temporal separation of the frequency components (Fig. 1a, Fig. S1). The beating frequency of the two beams is linearly correlated with the temporal delay between the two laser pulses. Using a two-dimensional galvo scanner, the combined laser beam is moved across the x and y dimensions of the sample to generate an image. This process is then repeated for a range of temporal delays, each of which produces a different wavenumber, ultimately producing a hyperspectral SRS image generated in a frame-by-frame manner. The spectral region surrounding the 2900 cm^-1^ wavenumber is typically referred to as the ‘C-H region’ and has a strong SRS signal. Biomolecules such as proteins and fatty acids, which contain many C-H bonds, show high Raman signal in this region. Importantly, SRS intensity scales linearly with molecular concentrations. The strong signal in the C-H region enables high fidelity SRS imaging with low optical powers that are compatible with live-cell imaging. Thus, this configuration can be used to acquire longitudinal images of live cells, resulting in data across four dimensions: space (x and y), wavenumber, and time. We set out to utilize SRS chemical imaging in the C-H region to measure fatty acid production in metabolically engineered strains of *E. coli*.

**Figure 1.**
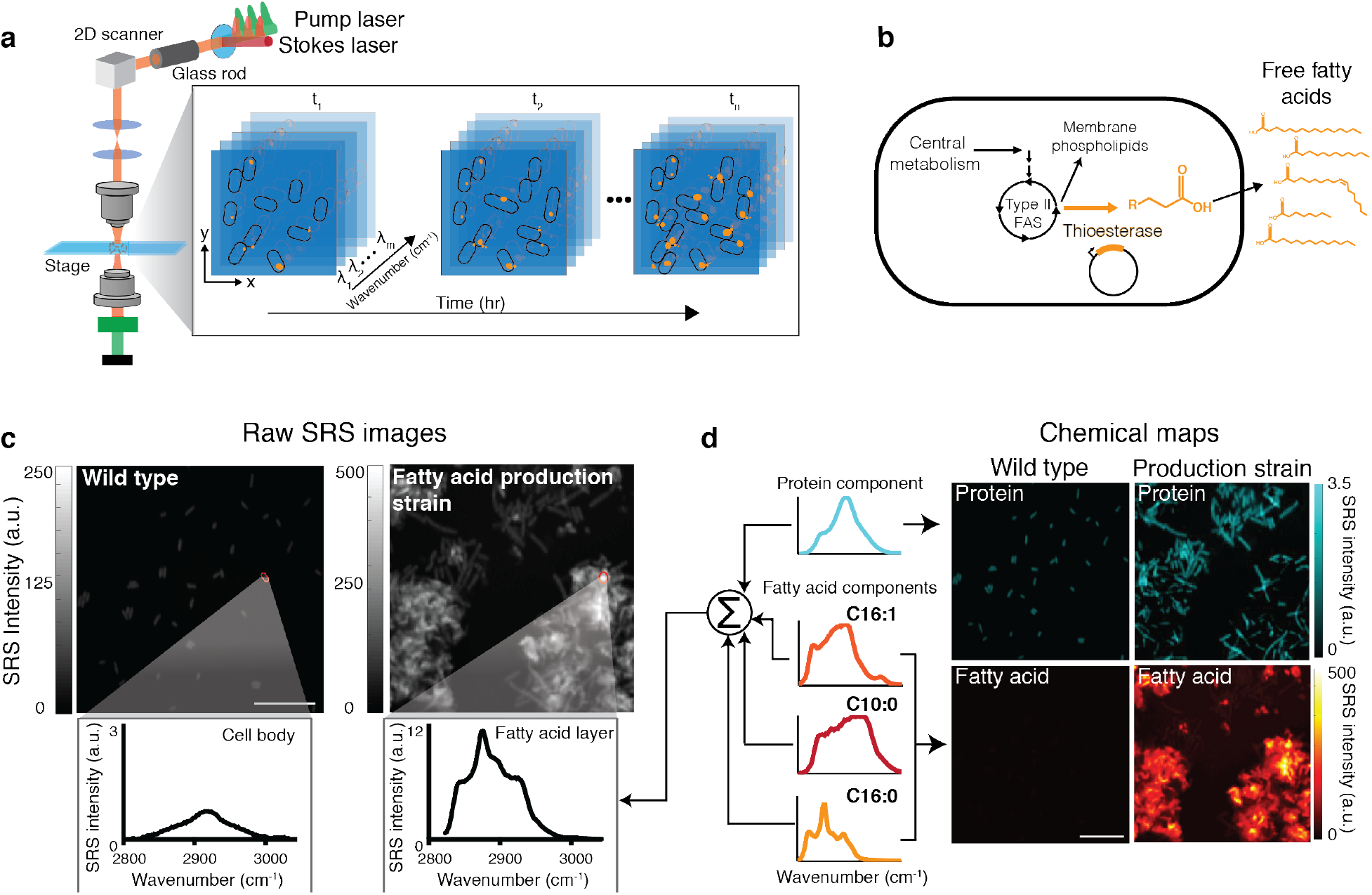
SRS imaging of *E. coli* production strains shows single-cell free fatty acid levels. **(a)** Schematic of the optical setup for SRS imaging to produce hyperspectral images using a Stokes and pump laser focused on a live sample. Hyperspectral SRS images contain three-dimensional data: x and y coordinates and wavenumber, which provides spectral information. Longitudinal SRS imaging adds a fourth dimension, time. **(b)** Schematic of free fatty acid production in *E. coli*. Expression of a cytosolic thioesterase results in free fatty acid accumulation through the type II fatty acid synthesis (FAS) pathway. Free fatty acids can vary in chain length and unsaturation, largely dictated by thioesterase specificity. **(c)** Representative raw SRS data from wild type *E. coli* and a strain overexpressing a cytosolic thioesterase (*Ab*TE*). The summation of Raman spectra at each pixel is shown. Representative regions are outlined in red with the corresponding Raman spectra shown below the image. Fatty acids and proteins emit strong Raman signals in the C–H region (∼2900 cm^-1^). Note that the y-axis scales are different; Fig. S3 shows them on the same scale. Scale bar, 10 μm. **(d)** Spectra at each pixel of the SRS image can be decomposed to generate chemical maps. Protein and fatty acid components are decomposed using spectral standards to produce chemical maps. Spectral standards shown in schematic are Bovine serum albumin (cyan), palmitoleic acid (C16:1, orange), capric acid (C10:0, red), and palmitic acid (C16:0, yellow). Protein and fatty acid chemical maps for both strains are shown. Scale bar, 10 μm.

Previous metabolic engineering efforts have focused on producing free fatty acids in *E. coli* using the native type II fatty acid synthesis pathway (14, 20, 34). Introducing a heterologously expressed acyl-acyl carrier protein (ACP) thioesterase can catalyze the formation and pooling of free fatty acids from elongating acyl hydrocarbon chains that would otherwise be incorporated into membrane phospholipids (35, 36) (Fig. 1b). We reasoned that SRS imaging could effectively capture fatty acid in production strains due to the C-H-rich carbon chains present in fatty acids. To test this hypothesis, we studied several production strains that were previously engineered to produce high quantities of free fatty acids (Tables 1 and 2). We first focused on the strain *Ab*TE*, which expresses an acyl-ACP thioesterase from *Acinetobacter baylyi*, carrying G17R/A165R mutations that improve enzymatic activity in *E. coli* (37). SRS images of *Ab*TE* show increased fatty acid production relative to the wild type strain, as evidenced by differences in both the chemical spectra and visible fatty acid droplets around the cells (Fig. 1c). Using spectral standards, SRS images can be decomposed into their major chemical components to produce chemical maps (Fig. 1d). We used standard spectra from pure protein (Bovine serum albumin, BSA), saturated fatty acids (C10:0 and C16:0), and unsaturated fatty acids (C16:1) to decompose the hyperspectral image (Fig. S2). To achieve this, we used a least absolute shrinkage and selection operator (LASSO) linear unmixing analysis to separate the hyperspectral image into its chemical components (Methods). This results in two dimensional chemical maps for protein and fatty acid components. Protein levels were comparable between wild type and *Ab*TE* strains, with slightly elevated levels in the engineered strain. In contrast, the fatty acid signal in the *Ab*TE* strain was significantly stronger than in wild type. Wild type cells contain membrane phospholipids, however these signals are much weaker than those recorded in the *Ab*TE* strain (Fig. S3). It should be noted that these strains were sampled from liquid culture, where free fatty acids are secreted and can aggregate in the media. As a consequence, the large fatty acid drops are not necessarily produced by the cells within the field of view, but could be an aggregate of fatty acid produced from many cells in the liquid culture. In subsequent studies we address this by growing cells on agarose pads to allow for affiliation of cells and the fatty acids they produce, however snapshots from liquid culture provide a view into the aggregate production.

**Table 1.** List of plasmids used in this study.

| Plasmid | Origin | Overexpressed operon | Resistance | Reference |
| --- | --- | --- | --- | --- |
| pSS200 | pMB1 | P<sub>trc</sub>-*Abte*:G17R/A165R | Amp<sup>R</sup> | Sarria *et al.* (37) |
| pBbA5c-‘tesA-vhb50-8fadR | p15a | P<sub>lacUV5</sub>-‘tesA-vhb50, P<sub>BAD</sub>-fadR | Cm<sup>R</sup> | Liu *et al.* (47) |
| pBbA5c-vhb50-8fadR | p15a | P<sub>lacUV5</sub>-vhb50, P<sub>BAD</sub>-fadR | Cm<sup>R</sup> | This study |
| pBbA5c-CpFatB1.2-M4-287 | p15a | P<sub>lacUV5</sub>-CpfatB1.2-M4-287 | Cm<sup>R</sup> | This study, mutant enzyme from Hernandez Lozada *et al.* (46) |
| pBbA5c-‘tesA-sfGFP-vhb50-8fadR | p15a | P<sub>lacUV5</sub>-‘tesA-sfGFP-vhb50, P<sub>BAD</sub>-fadR | Cm<sup>R</sup> | This study |
| pSS200-sfGFP | pMB1 | P<sub>trc</sub>-*Abte*:G17R/A165R-sfGFP | Amp<sup>R</sup> | This study |
| pBbE-ibpAB-k-mRFP1 | ColE1 | P<sub>ibpAB</sub>-mRFP1 | Kan<sup>R</sup> | This study, based on promoter from Ceroni *et al.* (53) |

**Table 2.** List of *E. coli* strains used in this study.

| Strain | Relevant genotype | Reference |
| --- | --- | --- |
| BW25113 (wild type) | F<sup>-</sup> Δ(araD-araB)567 ΔlacZ4787(::rrnB-3) λ<sup>-</sup> rph-1 Δ(rhaD-rhaB)568 hsdR514 | Baba *et al.* (71) |
| BW25113 Δ*fadE* | *E. coli* BW25113 Δ*fadE*, cured from Keio collection | Baba *et al.* (71) |
| MG1655 | F<sup>-</sup>, λ<sup>-</sup>, rph-1 | Blattner *et al.* (77) |
| *AbTE* * | *E. coli* MG1655; pSS200 | Sarria *et al.* (37) |
| ‘TesA-FV50 | *E. coli* BW25113 Δ*fadE*; pBbA5c-‘tesA-vhb50-8fadR | Liu *et al.* (47) |
| *AbTE* *-FV50 | *E. coli* MG1655; pBbA5c-vhb50-8fadR, pSS200 | This study |
| *CpFatB1* * | *E. coli* MG1655; pBbA5c-CpfatB1.2-M4-287 | This study |
| ‘TesA-FV50-sfGFP | *E. coli* BW25113 Δ*fadE*; pBbA5c-‘tesA-sfGFP-vhb50-8fadR | This study |
| *AbTE* *-sfGFP-FV50 | *E. coli* MG1655; pBbA5c-vhb50-8fadR, pSS200-sfGFP | This study |

### Characterization of enzymatic specificity, chain length distribution, and degree of unsaturation

Analytical chemistry methods such as GC-MS are typically employed to measure chemical production because they offer precise chemical specificity information. For fatty acid quantification, gas chromatography effectively separates fatty acid esters based on chain length and, along with mass/charge spectra, can specifically read out fatty acid ester chain length and unsaturated bonds. From a metabolic engineering perspective, quantification of a fatty acid production strain’s chain length distribution and level of unsaturation are critical. For biofuel purposes, chain length and termination chemistry can be tuned to mimic characteristics of fuel sources such as gasoline, diesel, or jet fuel (38). Alternatively, medium chain fatty acids (C8-C12) and their derivatives can be sources of many specialty chemicals (39). With these end point applications in mind, we sought to extend SRS imaging capabilities to capture the specific profiles of free fatty acid production strains.

Although pure fatty acids of different chain lengths have different spectra in the C-H region, they are too similar to accurately decompose using spectral unmixing with LASSO linear regression analysis. However, we expanded our analysis methodology to take advantage of spectral windows that correspond to CH_2_ or CH_3_ bonds, which are present in the 2832-2888 cm^-1^ and 2909-2967 cm^-1^ wavenumber regions, respectively (40). Since a saturated fatty acid has an increasing number of CH_2_ bonds as the chain length increases, but the terminal CH_3_ bond number is constant, we reasoned that the ratio of the CH_2_/CH_3_ spectral windows would scale with chain length (Fig. 2a). Using pure saturated fatty acid standards of variable chain length, we observed a nearly linear (R^2^ = 0.97) relationship between chain length and the ratio of CH_2_/CH_3_ area under the curve (Fig. 2b). We next tested whether we could use this relationship to estimate chain length production profiles.

**Figure 2.**
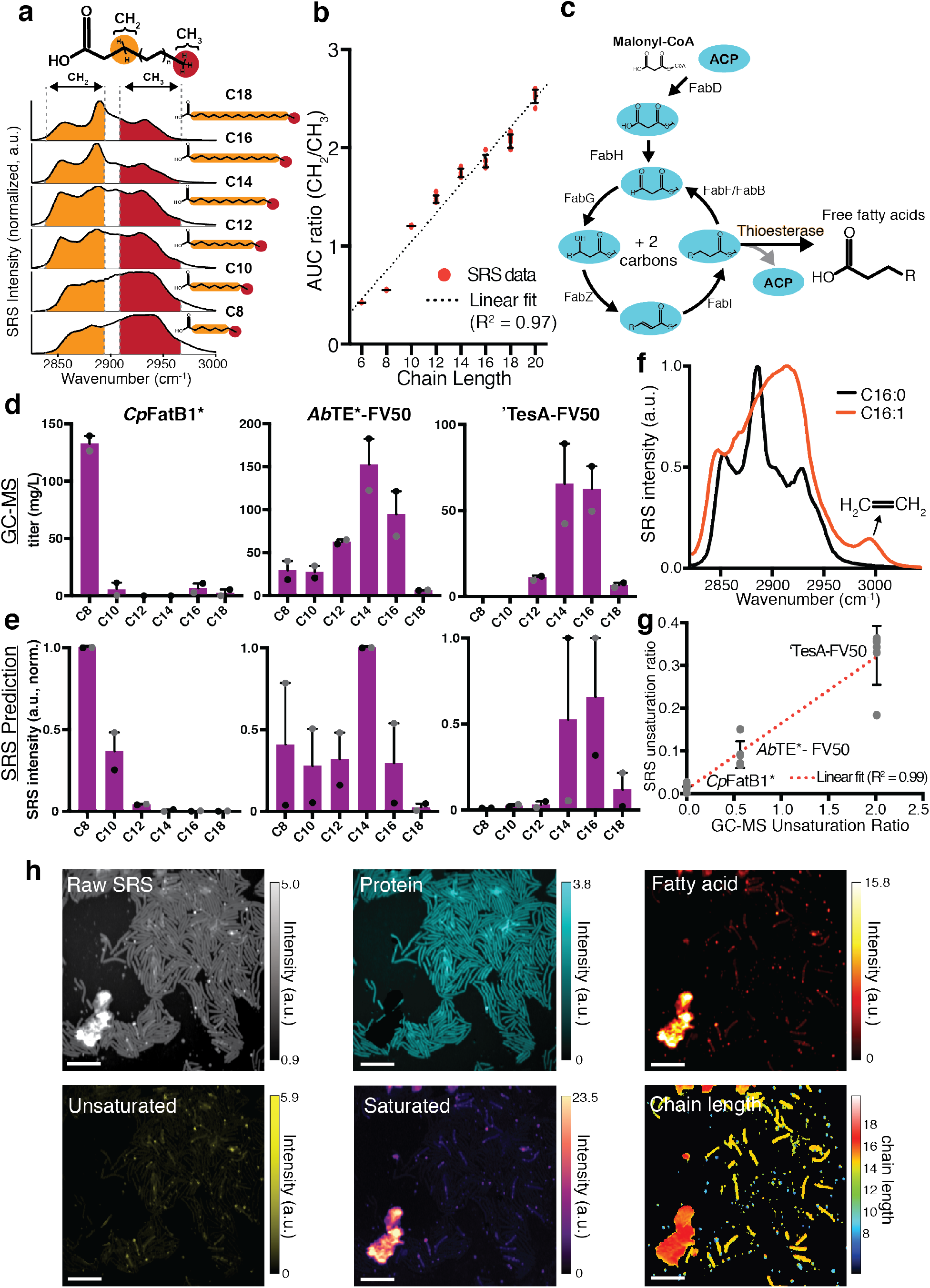
Chain length distribution prediction from different thioesterase enzymes. **(a)** The ratio of internal CH_2_ and terminal CH_3_ bonds within a fatty acid is a function of chain length. Raman spectra of pure fatty acid standards are shown for different chain lengths. Specific spectral windows correspond to each bond. **(b)** The ratio of area under the curve (AUC) of CH_2_/CH_3_ bonds scales approximately linearly with chain length. Error bars show standard deviation of n = 6 replicates. **(c)** Schematic of the type II fatty acid synthesis pathway in *E. coli*. Introduction of an acyl-ACP thioesterase pulls out elongating acyl-ACPs to form free fatty acids. Enzymatic specificity of the thioesterase largely determines the distribution of the fatty acid chain length profile. **(d)** Chain length distribution prediction with GC-MS compared to **(e)** SRS using CH_2_/CH_3_ ratio analysis (n = 2 biological replicates using 5 fields of view for each replicate, errors bars show standard error). Strains shown are: *Cp*FatB1*, *Ab*TE*-FV50, and ‘TesA-FV50 (Table 2). **(f)** SRS spectra of saturated and unsaturated fatty acid standards (C16:0, C16:1). The unique peak at ∼3000 cm^-1^ allows for spectral decomposition of unsaturation content. **(g)** Comparing GC-MS unsaturation ratio of produced free fatty acids to SRS production based on spectral analysis. Error bars show standard deviation from n = 5 fields of view for each strain. **(h)** Spectral decomposition and chain length prediction of *Ab*TE*-FV50 grown on an agarose pad. Scale bars, 10 μm.

In *E. coli*, fatty acid biosynthesis is carried out through a multistep, enzymatic Claisen reduction (41). The enzymatic components of type II fatty acid synthesis in *E. coli* are encoded as separate proteins, creating a pathway in which two carbons are added to an elongating acyl-ACP chain with each cycle (Fig. 2c). The number of cycles around this pathway before the elongating acyl chain is cleaved by an acyl-ACP thioesterase determines the resulting fatty acid chain length. The primary factor driving chain length is thought to be the enzymatic specificity of the heterologously expressed thioesterase (11, 42). Researchers have carried out numerous efforts to engineer specificity of acyl-ACP thioesterases in order to create desired chain length profiles (14, 37, 43–45). Several thioesterases have been shown previously to produce a range of free fatty acid chain length profiles. Three examples are *Cp*FatB1*, *Ab*TE*, and ‘TesA. The *Cp*FatB1* and *Ab*TE* thioesterases originate from *Cuphea palustris* and *A. baylyi*, respectively, and the “*” denotes mutants that were engineered to increase activity in *E. coli* (37, 46). ‘TesA is *E. coli*’s native thioesterase, where the “‘” denotes deletion of the leader sequence (35). Endogenously, TesA contains a leader sequence that localizes the enzyme to the periplasm; deleting the leader peptide sequence allows for interaction with cytosolic acyl-ACPs, enabling the production of free fatty acids (35) (Fig. S4).

To test our ability to estimate chain length distributions using imaging, we examined strains *Cp*FatB1*, *Ab*TE*-FV50, and ‘TesA-FV50, which each express a different thioesterase (Table 1, Table 2). Strains *Ab*TE*-FV50 and ‘TesA-FV50 additionally express heterologous *fadR* and *vhb50*, which have been shown to increase free fatty acid production (12, 47). FadR is a transcription factor that regulates many genes in the fatty acid synthesis pathway for increased free fatty acid titer when expressed alongside ‘TesA. Vhb50 is a *Vitreoscilla* hemoglobin that further increases fatty acid production by increasing oxygen uptake. We conducted an experiment in which each of the three strains were grown in liquid culture and thioesterase expression was induced for 24 hours to produce free fatty acids. Samples from each production culture were taken in parallel for GC-MS quantification and SRS hyperspectral imaging. As expected, GC-MS results show highly variable chain length distributions depending on the thioesterase expressed (Fig. 2d). *Cp*FatB1* primarily produces octanoic acid (C8:0). *Ab*TE*-FV50 produces a mix of medium- and long-chain saturated fatty acids with myristic acid (C14:0) as the largest component. Lastly, ‘TesA-FV50 produces long-chain fatty acids with large contributions from both myristic (C14:0) and palmitic acid (C16:0). Since each production strain has a unique chain length profile, they serve as an ideal group of strains to test our ability to predict chain length distributions with SRS imaging.

To implement chain length prediction, we first decomposed the spectra at each pixel into protein and representative fatty acid chemical maps (C10:0, C16:0, C16:1). The protein and unsaturated fatty acid maps were then subtracted from the raw SRS image to produce a hyperspectral SRS image of saturated fatty acids (Fig. S5), which can be used to estimate the average chain length at each pixel. We introduced a concentration weighting factor using the SRS spectral ensemble intensity at the same pixel. The SRS predicted chain length distributions closely matches the qualitative features of the GC-MS distributions (Fig. 2e). Importantly, the prediction captures whether the strain produces primarily medium- or long-chain fatty acids, or a mixture of both. In the case of ‘TesA-FV50, which produces primarily a mixture of C14 and C16, the SRS prediction results in either chain length largely dominating. This may stem from the binning needed during analysis to make a digital, even length prediction. For example, if a mixture of chain lengths is not spatially separated, a pixel prediction of 14.9 will result in a binary chain length prediction of all C14 (Methods). However, using several samples can correct for this type of issue, as seen in the average chain length prediction for ‘TesA-FV50.

To gauge unsaturation levels, we utilized the presence of the Raman peak at ∼3000 cm^-1^, which is unique to the C=CH_2_ bonds in unsaturated fatty acids (Fig. 2f). This peak serves as an identifier of unsaturation level and components from this fatty acid source can be unmixed with LASSO regression. To demonstrate our ability to predict unsaturation level from production strains, we tested the same three strains, which have different ratios of unsaturation to saturation (Fig. S4). The ratio of unsaturation from GC-MS data scales linearly with predicted unsaturated ratios from SRS images (Fig. 2g), giving an indication of the ability of this approach to predict the ratio of unsaturation. With the ability to calculate unsaturation level in addition to chain length distributions from SRS images, we cover many aspects of free fatty acid production that are important for metabolic engineers, bringing SRS hyperspectral imaging closer to a form of optical mass spectrometry.

We next applied our compositional analysis to *Ab*TE*-FV50 seeded and grown on agarose pads (Fig. 2h). Highly productive strains will secrete end-products, making it difficult to track the source of produced chemicals back to the cells that generated them. Therefore, sampling from liquid culture for imaging does not accurately provide production heterogeneity information. To ensure that free fatty acid production is tracked to the cells responsible for production, we first grew cells on agarose pads such that production could be localized to the region containing the cells. We observed a large aggregate of fatty acid outside the cells that is primarily composed of saturated, long chain fatty acids. This differs from interpretations of GC-MS quantification where it is assumed that long chain fatty acids remain within the cell (37). Additionally, single-cell chain length maps display a relatively homogenous makeup of chain lengths between individual cells, which is consistent with current understanding of the fatty acid synthesis pathway and thioesterase specificity (15). However, without single-cell resolution it would not be possible to distinguish between this scenario and one where chain length mixtures produced from bulk culture originate from distinct subpopulations that produce primarily one chain length each.

### Quantification of heterogeneity in fatty acid production strains

Given our ability to image production at the single-cell level, we asked whether our strains displayed production heterogeneity in the overall levels of fatty acid produced. Previous studies have reported sub-populations within production cultures that are less productive and lead to decreased overall performance of the population in a scaled up bioprocess (23, 48). Single-cell chemical imaging with SRS is uniquely suited to quantifying this phenomenon. We focused on strains *Ab*TE*-FV50 and ‘TesA-FV50 for agarose pad experiments because *Cp*FatB1* displayed poor growth in the agarose pad conditions.

We first quantified fatty acid production from *E. coli* microcolonies of the wild type and ‘TesA-FV50 production strain (Fig. 3a). Interestingly, ‘TesA-FV50 microcolonies exhibit a high level of colony-to-colony production variation. This intercolony heterogeneity is visible in the fatty acid chemical maps, with strains from the same original source exhibiting high and low producing microcolonies. One possible explanation for these differences in production is variable transcriptional regulation of key enzymes that are maintained through replication, leading to metabolic bottlenecks (7, 49). Alternatively, the ability to manage toxicity associated with production in the time frame following thioesterase induction may lead to divergent production outcomes (50).

**Figure 3.**
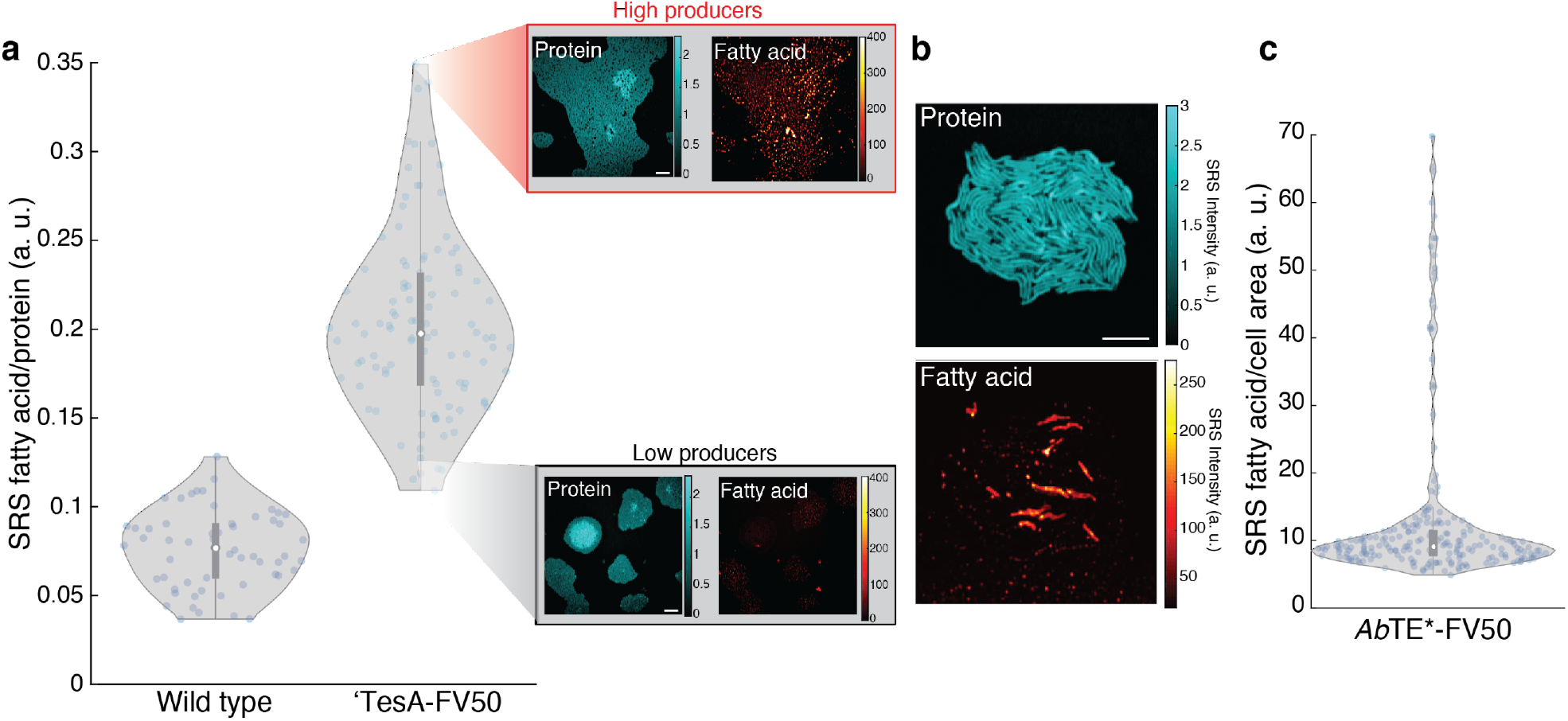
Inter- and intra-colony heterogeneity profiles of production strains. **(a)** Production from replicate ‘TesA-FV50 microcolonies (n = 105) are compared to wild type microcolonies (n = 56), revealing inter-colony production heterogeneity. Each data point represents fatty acid production from a single microcolony. Protein and fatty acid chemical maps are shown for representative high and low producing microcolonies. Scale bar, 10 μm. **(b)** Representative protein and fatty acid chemical maps are shown for a microcolony of the production strain *Ab*TE*-FV50. **(c)** Intra-colony production is quantified for single cells within the microcolony (n = 213) (Fig. S6). Each data point represents a single cells’ production. Scale bar, 10 μm. Box plot overlays contain median (white circle), first and third quartiles (gray box) and 1.5x interquartile range (thin gray line) for each distribution.

We also examined production heterogeneity in the fatty acid production strain, *Ab*TE*-FV50. Strikingly, we observed a very different type of production variation in this strain (Fig. 3b). Unlike the intercolony heterogeneity in ‘TesA-FV50, the *Ab*TE*-FV50 strain has high heterogeneity between cells in a single microcolony. We used the protein channel to segment the image into single cells for analysis (Fig. S6) and quantified single-cell production (Fig. 3c). Our quantification indicates that in this strain a small percentage of cells produce the vast majority of fatty acids. This result is consistent across many fields of view within the microscopy images, suggesting that it is a general feature of this production strain (Fig. S7).

### Longitudinal SRS imaging of fatty acid production during growth of colonies

Understanding the dynamics of chemical production with single-cell resolution can provide key insights into the emergence of heterogeneity, production bottlenecks, and can guide engineering strategies to maximize metabolic flux. To that end, we sought to adapt the SRS system for longitudinal imaging. While SRS imaging of living cells has been reported (26, 51, 52), its application to chemical production over long periods of growth has not been demonstrated. Previous work from Wakisaka, *et al*. achieved video rate SRS for short periods of time by reducing spectral acquisitions to four points in the C-H region (26). For metabolic engineering applications, however, spectral fidelity and time scales on the order of bioprocesses would provide a more useful form of longitudinal imaging. Therefore, we sought to develop parameters amenable to longitudinal imaging without loss of spectral information. We installed an incubator on the microscope stage and grew live cells on agarose pads for at least 16 hours at 31°C. First, we tested whether the routine laser powers we used for endpoint SRS imaging were damaging to live cells (75 mW for 1040 nm Stokes and 15 mW for 800 nm pump at the sample). At the beginning of longitudinal imaging, we captured a bright field transmission image and measured a hyperspectral SRS image in one field of view (Fig. S8a-b). After 16 hours of incubation, cells that were previously exposed to SRS imaging did not duplicate, nor did they produce significant levels of fatty acids. In contrast, cells in a region in the immediate vicinity that had not been exposed to imaging grew into a dense microcolony and produced fatty acid droplets (Fig. S8c-d). Although the laser exposure did not induce visible cell damage, the photodamage altered cell growth, indicating that these laser powers were too high.

To optimize the imaging conditions to reduce phototoxicity, we performed the same live-cell experiment with lower laser powers. We obtained normal cell growth when we reduced the Stokes power from 75 to 25 mW, while the pump laser at 800 nm was kept as 15 mW (Fig. S9). To illustrate growth and fatty acid production, we measured transmission and SRS images for the same field of view after 3 and 5 hours of incubation, seeing clear evidence of replication even after SRS imaging. We took a final wide-field image at 6 hours, which showed that cells continued to replicate normally, demonstrating that these laser power parameters permit growth. To further probe how these imaging conditions impact cells, we utilized a stress-responsive promoter, P_ibpAB_, to drive expression of mRFP1 (Fig. S10a). P_ibpAB_ is driven by the heat shock σ-factor (σ^32^) and is upregulated in response to stress (53). We first exposed cells to the 25 mW / 15 mW laser intensities describe above and compared promoter activity to cells that received no SRS exposure (Fig. S10b). Although these cells were able to grow, RFP expression indicates that intracellular stress was significantly upregulated in response to SRS exposure. To lower laser exposure further, we increased the step size of each laser scan from 150 nm to 230 nm, corresponding to fewer pixels per image. With the increased step size, RFP expression showed no significant difference relative to the cells that received no laser exposure (Fig. S10b). Therefore, we concluded that using both reduced laser powers and increased step size can allow for longitudinal SRS imaging.

With these optimized imaging conditions, we first tracked fatty acid production within the strain ‘TesA-FV50. In line with heterogeneity patterns we originally observed in this strain (Fig. 3a), the production trajectories varied across microcolonies (Fig. 4a-b). In one example, fatty acid signals increased in cells starting ∼12 hours after thioesterase induction (Fig. 4a). After the microcolony reached a high cell density on the agarose pad, we observed significant accumulation of fatty acids. In contrast, a second microcolony of the same strain produced only low levels of fatty acid (Fig. 4b). For comparison, we also tracked the growth and fatty acid production of wild type cells under the same conditions, observing only low levels of fatty acid production (Fig. S11). Time-lapse wide-field transmission images for the wild type strain show that cells under SRS laser exposure grew well during the entire experiment period and at levels comparable to those regions not exposed to imaging, reaffirming that these conditions are non-toxic (Movie S1). We quantified fatty acid and protein levels of each microcolony and the wild type strain. Protein levels in each strain increased at comparable rates (Fig. 4c). Fatty acid levels in the wild type colony increased modestly while the high-producing ‘TesA-FV50 microcolony fatty acid levels increased dramatically (Fig. 4d). The low-producing ‘TesA-FV50 microcolony produced fatty acids at levels comparable to wild type.

**Figure 4.**
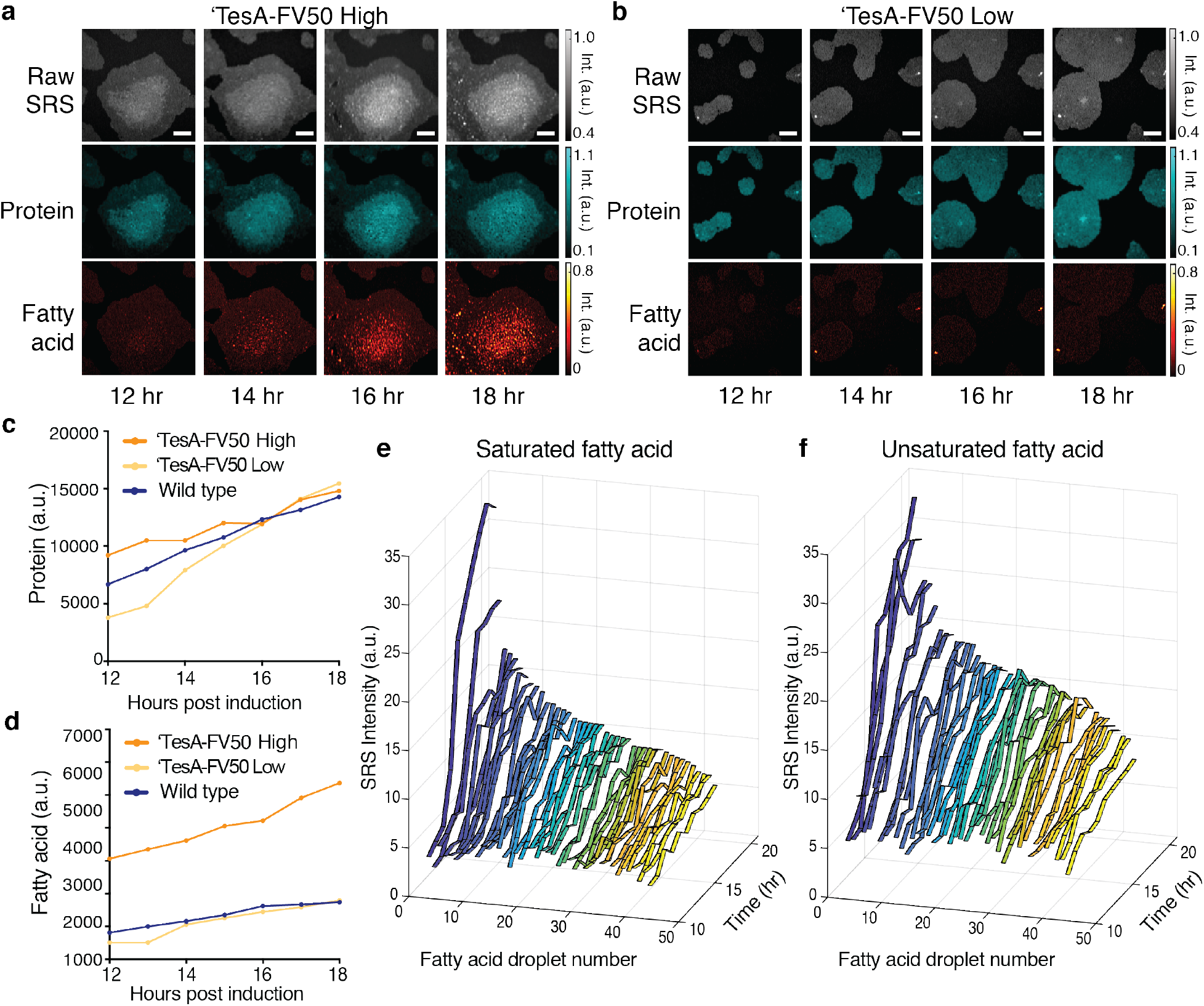
Longitudinal SRS imaging of production dynamics. Time-lapse images of **(a)** a ‘TesA-FV50 high producing microcolony and **(b)** a ‘TesA-FV50 low producing microcolony under the same conditions, shown with the raw SRS images (spectral summation of the SRS image stack) and chemical maps corresponding to protein and fatty acid content. Scale bars, 10 μm. Quantification of **(c)** protein and **(d)** fatty acid over time from the microcolonies in (a-b and Fig. S11). **(e)** Saturated and **(f)** unsaturation content of individual droplets from the ‘TesA-FV50 high microcolony shown in (a). Droplets are numbered in order of their final fatty acid levels and numbers are consistent between (e) and (f).

The activity in the high-producing ‘TesA-FV50 microcolony is in line with known regulation patterns in *E. coli* fatty acid synthesis. When high cell density is reached in wild type *E. coli*, the pathway is inhibited by a buildup of acyl-ACPs. This mechanism is reported to act through direct inhibitory interactions with key enzymes within the pathway, such as acetyl-CoA carboxylase, FabH, and FabI (54, 55). Additionally, acyl-ACP or acyl-CoA responsive transcription factors, FadR and FabR, respectively, act to regulate transcriptional responses that control fatty acid synthesis (56, 57). In the presence of a cytosolic thioesterase, as in the ‘TesA-FV50 strain, this inhibition is released through the conversion of accumulated acyl-ACPs to free fatty acids. However, thioesterase expression is induced starting at t = 0 hr, and significant accumulation of fatty acid does not happen until the microcolony is well established. Even with the ‘TesA thioesterase highly expressed, phospholipid metabolism may dominate metabolic flux through the fatty acid synthesis pathway until sufficient density is reached to suppress incorporation of acyl-ACPs into phospholipids. A recent study from Noga *et al*. uncovered a post-translational mechanism that modulates phospholipid biosynthesis through PlsB acyltransferase and ppGpp, which may explain the delay in free fatty acid accumulation (58).

Additionally, we measured the dynamics of the *Ab*TE*-FV50 fatty acid production strain at the microcolony level, which produces a variety of medium- and long-chain fatty acids (Fig. S4), with significant heterogeneity in production among cells (Fig. 3b-c). We again observed fatty acid production over time, with similar delays in fatty acid accumulation despite thioesterase induction at t = 0 hr (Fig. S12a). In this strain, a few cells within the microcolony produce large amounts of fatty acid. The production dynamics for these few cells are similar to fatty acid production within the ‘TesA-FV50 strain, but the remainder of cells exhibit at low levels of production for the duration of imaging.

To further understand the dynamics of fatty acid production, we tracked the composition of individual droplets from the high producing ‘TesA-FV50 microcolony and high producing cells from the *Ab*TE*-FV50 microcolony. Both saturated and unsaturated fatty acid levels increase similarly within the droplets of the ‘TesA-FV50 strain (Fig. 4e-f). Interestingly, the high producing cells from the *Ab*TE*-FV50 strain initially produce saturated fatty acids, but saturated fatty acid levels plateau in a subset of cells as the incubation continues (Fig. S12b). Alternatively, unsaturated fatty acid production continues to increase for the duration of the experiment (Fig. S12c). Additionally, we analyzed the chain length composition for both strains longitudinally (Fig. S13a-b). Droplets from ‘TesA-FV50 ranged from C14-C16 in length, which is in line with bulk culture production. Chain lengths for *Ab*TE*-FV50 high producer cells displayed high fluctuations over time and range from C7-C14, which is shorter than expected in comparison with bulk culture data. We believe the fluctuations and low chain length predictions stem from a decreased signal-to-noise ratio using our low power parameters for longitudinal imaging. When the signal-to-noise ratio is increased for stronger SRS signals, such as for the large extracellular droplet within the *Ab*TE*-FV50 microcolony, the chain length prediction increases to a range between C12-C14, which more closely matches bulk culture data (Fig. S12a, Fig. S13b).

### Single cell growth-production relationship

Next, we asked whether cell-to-cell differences in fatty acid production correlate with differences in growth rates between cells. Production of a heterologous product is often associated with changes in cell physiology due to the consumption of resources and intermediate or end-product associated toxicities (59–61). Consequently, we asked whether growth rate is inversely correlated with fatty acid production. For this analysis, we focused on the *Ab*TE*-FV50 strain because it exhibits significant intracolony heterogeneity. At the bulk culture level, we do not observe a decrease in growth when production is induced through *Ab*TE* expression (Fig. S14a-b). However, bulk culture measurements do not rule out slow growth of a high-producing subpopulation. To understand whether there exists a growth tradeoff in the high producer subpopulation, we measured growth at the single-cell level. Although we can resolve single cells using the longitudinal SRS conditions, the lowered resolution needed to avoid phototoxicity hinders single-cell segmentation to quantitatively probe growth at many time points. To avoid these limitations, we used a combination of time-lapse, phase contrast microscopy followed by endpoint SRS imaging (Fig. 5a). Using the high-resolution phase contrast images, we then segmented and quantified single-cell growth rates using an automated segmentation pipeline for microcolonies (62). Pairing growth quantification with endpoint SRS, we tracked the growth trajectories and lineages of single cells within the microcolony to their fatty acid production. Spectral decomposition of the endpoint SRS image allows the high fatty acid cells to be identified, along with other chemical composition information (Fig. 5b). Growth of the high producer cells in the microcolony, measured as cell length over time, did not correlate with lower growth rates (Fig. 5c, Fig. S15, Movies S2-4). We binned cells into two groups, low and high fatty acid producers, where we defined high producers as those with production in the top 15% of single cells in the distribution (Fig. S16). Examining the growth rates of each cell near the endpoint (16 hr) and earlier in the time course (8 hr) shows that growth rate is not significantly different between the high and low producers.

**Figure 5.**
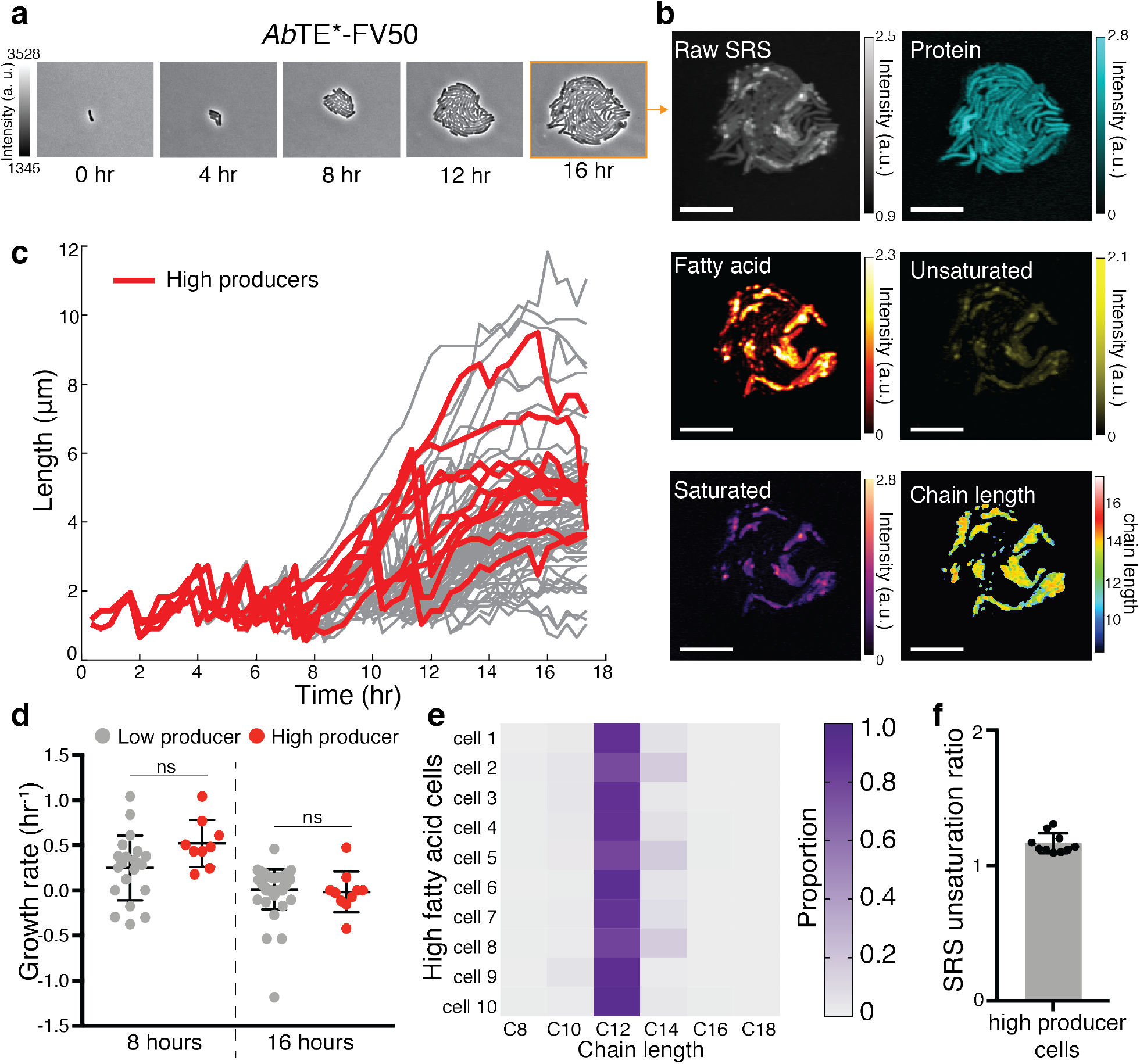
Single cell growth-production relationship. **(a)** Time-lapse phase contrast imaging of an *Ab*TE*-FV50 microcolony followed by **(b)** endpoint SRS imaging and spectral decomposition. **(c)** Single-cell lengths as a function of time within the microcolony shown in (a-b), with high producer trajectories highlighted in red (n = 68 cells). Sharp decreases in length mark a cell division. High producers are defined as the top 15% of producer cells (Fig. S16). **(d)** Growth rate comparisons of high and low producer trajectories at 8 and 16 hours (p = 0.0507 and p = 0.714, respectively; two tailed unpaired t-test). Growth rate is calculated from cell length data in (c) (see methods). **(e)** Saturated chain length prediction of high producer cells. **(f)** Unsaturation ratio (unsaturated/saturated) of high producer cells.

Given our ability to decompose the fatty acid signal into unsaturated and chain length components, we also analyzed the top 10 producer cells’ composition to gain further insight into the high fatty acid phenotype in this strain. In contrast with GC-MS measurements sampled from bulk culture, each cell is enriched with lauric acid (C12:0) relative to other saturated fatty acid chain lengths (Fig. 5e). Additionally, the unsaturation ratio of the top producers is significantly increased in high producer cells relative to bulk culture sampling (Fig. 5f, Fig. 2g). The decreased levels of myristic acid (C14:0) and palmitic acid (C16:0) present in the high fatty acid cells relative to bulk culture may be related to unsaturated fatty acid biosynthesis. In *E. coli* fatty acid synthesis, double bonds in the carbon tail of elongating fatty acids are formed specifically when the carbon chain has reached decanoyl-ACP (C10), followed by further elongation to C12:1, C14:1, or C16:1 (63). It is possible that chain lengths that would have otherwise reached C14:0 and C16:0 are instead unsaturated.

## Discussion

Chemical imaging can play a key role in the strain engineering process. Current quantification techniques rely either on methods like GC-MS, which are chemically-specific but where information about individual cells and their dynamics are lost, or on fluorescent reporters or dyes, which are indirect readouts and can be difficult to engineer or limited in their specificity. SRS imaging has the potential to dramatically improve this process by providing key insights into chemical production at the single-cell level. Thus, methods that were previously only accessible with single-cell readouts, such as directed evolution or cell-sorting approaches are in principle possible with SRS imaging. Further, the ability to track production changes over time can provide insight into the emergence of production heterogeneity and, ultimately, guide strategies to avoid low producers in the population. The landscape for strain engineering is expanding rapidly, with systems biology approaches to enzyme engineering and novel technologies for quantifying production offering great promise for improving designs. In this study we focus on fatty acid synthesis, which is an important pathway that can be engineered to produce a diversity of valuable chemicals. Development of this pathway towards near theoretical yields will be important to replace many industrial chemicals with sustainable bio-based alternatives (5).

Here, we examined free fatty acid production strains of *E. coli* using SRS and demonstrated that hyperspectral imaging allows for image decomposition into major chemical components, with the ability to distinguish cells from their chemical product. By incorporating additional analysis, we also introduce an approach that can estimate chain length distribution and unsaturation degree, increasing the amount of information that can be extracted from SRS hyperspectral images. These advances can enable a metabolic engineer to examine fatty acid production strains using SRS imaging while maintaining important chemical specificity data. The ability to gauge enzyme specificity through imaging opens the possibility of screening mutant enzyme libraries in a high throughput fashion to select for optimal free fatty acid profiles.

Visualizing chemical production at the single-cell level reveals important information that would otherwise be obscured by bulk culture quantification methods. We demonstrate this by examining production heterogeneity among different engineered strains, observing both intra-and inter-colony differences in production within microcolonies. These results provoke fundamental questions about the mechanisms leading to cellular heterogeneity, and also suggest that engineering strategies that eliminate low-producers could improve yields. For example, it may be possible to gradually enhance the overall production levels of a strain of engineered *E. coli* through multiple cycles of growth and dilution, with a step that removes low-producers at the end of each cycle.

Furthermore, we established parameters that allow us to extend SRS imaging for longitudinal studies in live cells. Unlike previous phototoxicity studies focusing on acute responses like membrane blebbing (64, 65), we directly observe long-term cell functions including cell replication, free fatty acid synthesis, and the absence of induction of stress response. SRS imaging has been used to probe metabolic heterogeneity in live cells previously in an elegant study by Wakisaka *et al*. (26), and we extend these results in several critical ways. In our experiments we track the same cells over multiple hours, rather than sampling new cells from liquid culture at each timepoint. In addition, we use *E. coli* for our study while Wakisaka *et al*. use the alga *Euglena gracilis. E. coli* are highly amenable to metabolic engineering, but their small size makes both imaging and analysis more challenging (*E. coli* are 1-2 μm in length while *E. gracillis* are 35-50 μm (66)). Thus, our results significantly extend prior findings, offering longitudinal imaging of a highly relevant engineered species. We envision production tracking at the single-cell level will be valuable for metabolic engineering studies by establishing how and when heterogeneity emerges. To quantify single-cell properties such as growth rate, however, higher resolution longitudinal imaging is needed to achieve time lapse data that can be processed with segmentation algorithms. Further development focused on mitigating phototoxicity without decreasing resolution may be able to overcome this challenge in the future.

As we demonstrate, a hybrid approach using phase contrast imaging and endpoint SRS microscopy allows for fundamental questions to be examined, such as the growth-production tradeoff. Interestingly, in the *Ab*TE*-FV50 strain that we studied using this hybrid approach, we observed no tradeoff between growth and production. This information, along with insights into the composition of the high fatty acid cells, can lead to novel hypotheses of the underlying cause of intracolony heterogeneity in this strain. These results underpin the utility of examining single-cell characteristics to increase performance of a given strain. For example, recent approaches to increase bioproduction involving dynamic regulation, either through transcriptional feedback circuits or optogenetic regulation, show promise to increase strain efficiency (67, 68). Imaging single-cell production dynamics in these strains could increase our understanding of how feedback systems can be used in the context of metabolic engineering. Together with synthetic biology methods, our system has the potential to answer fundamental questions relating to the production of biosynthetic targets at the single-cell level. Further, because SRS imaging does not require engineered biosensors, it has the potential to serve as a widely useful platform to boost the pace of strain engineering for a broad range of metabolites.

Moving forward, it will be important to understanding the connection between production at the single-cell level and bulk culture output. Imaging fields of view sampled from bulk culture can potentially lead to biased overall titer prediction, especially if the product is not soluble in water. Alternatively, studying microcolonies grown on agarose pads is ideal for imaging but not necessarily predictive of bulk culture behaviors. For example, nutrient mixing, population selection, and secretion may differ between the two-dimensional growth conditions and a well-stirred liquid culture. Additionally, SRS has sensitivity limits significantly higher than mass spectrometry (69) and thus requires a product to be produced at sufficient quantities before SRS can be used to guide further engineering. Given these limitations, we envision that SRS studies will be most useful for strain optimization rather than enzyme or pathway discovery.

SRS imaging in different spectral regions, such as the fingerprint region (400-1800 cm^-1^), can be adapted to study strains producing non-fatty acid derived chemicals of interest, such as terpenes, to expand the scope of SRS imaging in metabolic engineering (29). In addition, because the approach is label-free it does not require biosensors with fluorescent reporter readouts, making it amenable to quantification of production in organisms that are recalcitrant to genetic modification.

Moreover, instrumentation advancements can enable SRS guided single-cell screening, such as SRS-based cell sorting, which has been demonstrated recently for cell phenotyping (70). The throughput we achieve in this study is limited by spectral tuning of the motorized delay stage and time spent manually focusing on samples. In future work, applying the ultrafast spectral tuning SRS system from Lin *et al*. (29), along with integrated autofocusing could drastically increase throughput. Much like the utility of fluorescence activated cell sorting in synthetic biology applications, we envision that SRS-based cell sorting could increase the throughput of strain screening and enable directed evolution based on chemical production. This work acts as a jumping off point for SRS imaging in metabolic engineering to aid in the development of more efficient strains for renewable chemical production.

## Methods

### Bacterial strains and plasmids

Plasmid and strain information are listed in Tables 1 and 2. The pBbA5c-’tesA-vhb50-8fadR plasmid was a gift from Dr. Fuzhong Zhang. The BW25113 Δ*fadE* strain is from the Keio collection (71), and we used the FLP recombination protocol from Datsenko and Wanner to cure the *kan*^*R*^ cassette from the genome (72). We used golden gate cloning (73) to create the pBbA5c-vhb50-8fadR plasmid by deleting the coding sequence of ‘*tesA* from pBbA5c-’tesA-vhb50-8fadR. The pBbA5c-CpFatB1.2-M4-287 plasmid was also constructed using golden gate cloning, with the pBbA5c backbone amplified from the BglBrick plasmid library (74) and the coding sequence of *Cp*FatB1.2-M4-287 derived from Hernández Lozada *et al*. (46) and synthetized by Twist Biosciences (South San Francisco, CA). pSS200 was a gift from Dr. Pamela Peralta-Yahya. pBbE-ibpAB-mRFP1 was constructed using the pBbE5k BglBrick backbone (74) with the promoter region of the genomic *ibpAB* operon as in Ceroni *et al* (53). We constructed pBbA5c-’tesA-sfGFP-vhb50-8fadR and pSS200-sfGFP using golden gate cloning with pBbA5c-’tesA-vhb50-8fadR and pSS200 as backbones, respectively, along with an sfGFP coding sequence containing a flexible GS linker to insert in frame with each thioesterase.

### Growth and induction of fatty acid production strains

For fatty acid production experiments, pre-cultures were grown overnight in LB media and used to inoculate 3 mL M9 minimal media (M9 salts, 2mM MgSO_4_, 100 μM CaCl_2_) with 2% glucose and grown at 37°C with 200 rpm shaking. Antibiotics were added to the media where necessary for plasmid maintenance according to resistances in Table 1 (100 μg/ml for carbenicillin and 25 μg/ml for chloramphenicol). The cultures were allowed to grow until approximately OD_600_ = 0.6 before thioesterase expression was induced with IPTG. Induction levels were 500 μM for ‘TesA-FV50 and 50 μM for *Ab*TE*, *Ab*TE*-FV50, and *Cp*FatB1*. For imaging from liquid cultures, cells were grown for 24 hours after IPTG induction and then 3 μL of sample was taken for imaging. Samples from liquid culture were placed on 3% agarose pads (Promega) containing M9 minimal media and sandwiched between glass coverslips to immobilize the cells for imaging. Samples from liquid culture were allowed to dry on the agarose pads for ∼15 minutes prior to imaging. For longitudinal imaging, production heterogeneity experiments, and phase contrast imaging, once cells reached OD_600_ = 0.6 in liquid culture, the sample was placed on a 3% low melting point agarose pad containing M9 minimal media with 2% glucose, IPTG as specified above, and appropriate antibiotics for plasmid maintenance, as detailed in Table 1. Microcolonies were imaged after 18 hours of growth on the agarose pads at 31°C.

For the chain length distribution prediction, cultures were induced with IPTG in liquid cultures for 24 hours. At the 24 hour timepoint, we took 3 μL of sample for imaging and another sample of the culture was taken for GC-MS analysis to allow direct comparison of the same culture. Five fields of view were imaged for each culture.

### Fatty acid derivatization and quantification with GC-MS

Samples for GC-MS quantification were taken at 24 hours post IPTG induction. 400 μL of vortexed culture was taken for fatty acid extraction and derivatization into fatty acid methyl esters as described by Sarria *et al*. (37) with the following minor modifications: Internal standards of nonanoic acid (C9) and pentadecanoic acid (C15) were added to the 400 μL sample at final concentrations of 88.8 mg/L each and vortexed for 5 sec. The following was then added to the sample for fatty acid extraction and vortexed for 30 sec: 50 μL 10% NaCl, 50 μL glacial acetic acid, and 200 μL ethyl acetate. The sample was then centrifuged at 12,000 g for 10 mins. After centrifugation, 100 μL of the ethyl acetate layer was mixed with 900 μL of a 30:1 mixture of methanol:HCl (12N) in a 2 mL microcentrifuge tube. The solution was vortexed for 30 sec followed by an incubation at 50°C for 60 mins for methyl ester derivatization. Once cooled to room temperature, 500 μL hexanes and 500 μL water were added to the 2 mL microcentrifuge tube, vortexed for 10 sec, and allowed to settle. 250 μL of the hexane layer was mixed with 250 μL ethyl acetate in a GC-MS vial for quantification.

The samples were analyzed with an Agilent 6890N/Agilent 5973 MS detector using a DB-5MS column. The inlet temperature was set to 300°C with flow at 4 mL/min. The oven heating program was initially set to 70°C for 1 min, followed by a ramp to 290°C at 30°C/min, and a final hold at 290°C for 1 min. GLC-20 and GLC-30 FAME standard mixes (Sigma) were tested using this protocol to ensure proper capture of all chain lengths and to gauge retention times. Internal standards were used for quantification, with chain lengths C8-C12 quantified with the nonanoic acid internal standard and C14-C18 quantified with the pentadecanoic internal standard.

### Optical setup

The SRS setup was driven by an 80 MHz femtosecond laser (Insight Deepsee+, Spectra Physics, USA) with two synchronized outputs. One output was fixed at 1040 nm with a pulse duration of ∼150 fs, while the other was tunable from 680 - 1300 nm with ∼120 fs pulse width. We used the 1040 nm beam as the Stokes and was modulated by an acousto-optical modulator (522c, Isomet, USA) at 2.5 MHz. We set the tunable output to 798 nm to excite the C-H region and spatially combined it with the Stokes by a dichroic mirror. Six 15 cm SF-57 glass rods were used to linearly chirp the femtosecond pulses to ∼ 2 ps. Five of the rods were placed on the common path while one was placed on the Stokes path to parallelize the degree of chirping considering its longer wavelength. A motorized delay stage was used to scan the temporal delay between two pulses to tune the excitation frequency. The combined beams were sent to a pair of two-dimensional galvo scanners (GVSM002, Thorlabs, USA) to perform laser scanning imaging. We used a 40X oil-immersion objective (RMS40X-PFO, Olympus, Japan) to focus the laser onto the sample. Powers on the sample were 15 mW for pump and 75 mW (or 25 mW for longitudinal imaging) for Stokes.

A home-built resonant amplifier photodiode collects and amplifies the stimulated Raman loss signal at the modulation frequency. We used a lock-in amplifier (UHFLI, Zurich Instruments, Switzerland) to extract the signal and send it to a data collection card (PCIe-6363, National Instruments, USA). We note that all elements described here are commercially available with the exception of the photodiode, which has been previously reported (75). Custom LabView (National Instruments, USA) software was used to synchronize the galvo scan with the delay line scan to obtain a hyperspectral SRS image stack in a frame-by-frame manner.

### Chemical map processing with LASSO

To obtain concentration maps for chemicals, we perform linear unmixing on the raw hyperspectral image stack. Assuming the number of pure components as *K* and the dimensions of a hyperspectral image as *N*_*x*_, *N*_*y*_, *N*_*λ*_, the unmixing model can be written as:

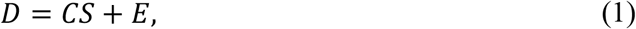

where 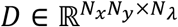 is the raw data reshaped as a two dimensional matrix in raster order, 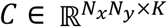 is the collection of concentration maps, 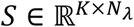 contains SRS spectra of all the components, while *E* is the residual term with error and noise. Given the prior knowledge of spectra for all the pure components, the task is reduced to generating chemical maps *C* via least square fitting. To avoid crosstalk between spectrally overlapped components, we add a *L*1 norm sparsity constraint by observing that at each spatial position, a few components dominate the contribution. The solution for *C* is found in a pixel-by-pixel manner by solving for the following optimization problem known as the least absolute shrinkage and selection operator (LASSO):

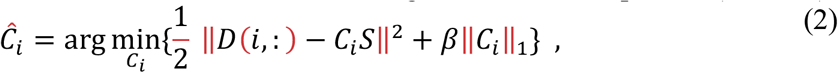

where *i* represents a specific pixel in the hyperspectral image, Ĉ*i* stands for the estimated concentrations for all components at pixel *i*, and *β* is a hyperparameter controlling the level of *L*1 norm regularization at each pixel.

For each imaging experiment, we measured spectra of pure chemical standards for analysis. Specifically, we input the spectra from the following pure components to perform linear unmixing: We use BSA as the protein standard, palmitic acid (C16:0) and capric acid (C10:0) as representative saturated fatty acids, and palmitoleic acid (C16:1) as an unsaturated fatty acid standard. All standards were sourced from Sigma Aldrich, USA.

### Chain length and unsaturation prediction

To predict chain length distribution, we first processed images with linear unmixing as described above. However, this analysis outputs two-dimensional chemical maps whereas a three-dimensional hyperspectral image is needed for chain length prediction. We created a hyperspectral, saturated fatty acid map by subtracting the protein and unsaturated fatty acid components from the original background-subtracted hyperspectral image (Fig. S5). We then calculated the area under the curve ratio of CH_2_ to CH_3_ for each pixel, using 2832 to 2888 nm for CH_2_ and 2909 to 2967 nm for CH_3_.

We used the linear relationship of ratio to chain length produced from standards (C6-C20, Sigma Aldrich, USA) to calculate a predicted chain length for each pixel. This prediction was then multiplied by a concentration weighting factor that corresponds to the SRS spectral summation at the same pixel. Thus, if the raw SRS signal from a region is low then its weight in the overall prediction is also low relative to pixels with strong SRS signal. All pixels’ in a field of view concentration-weighted chain lengths were compiled to create the fatty acid chain length distribution. To calculate the unsaturation ratio, the sum of the C16:1 chemical map generated through linear unmixing was divided by the sum of the hyperspectral saturated chemical map. For the tracking of fatty acids production and composition dynamics (Fig. 4e-f, Fig. S12, Fig. S13), we manually segmented significant fatty acid droplets using the fatty acid concentration map in the last time stamp. Each droplet was manually traced and segmented frame-by-frame in all earlier time stamps until no fatty acid was found (Movies S5-6).

### Single cell segmentation

Segmentation of single cells within SRS images was implemented in two steps. The protein segmentation map was first sent to CellProfiler to generate an initial segmentation (76). A customized pipeline was used for the analysis, including illumination correction, background subtraction, and edge enhancements based on the Laplacian of the Gaussian. Then a custom Matlab program was used to manually correct errors in the automated segmentation analysis using the raw SRS and protein chemical maps as a guide. When SRS images are segmented, we normalize the fatty acid channel by cell area instead of the protein channel. This normalization more accurately represents the single cell production, whereas the protein channel normalization at the microcolony level accounts for cells growing on top of each other. Since the primary source of heterogeneity in the *Ab*TE*-FV50 is at the single-cell level, we utilize the fatty acid intensity normalized to cell area metric. Alternatively, heterogeneity seen in the ‘TesA-FV50 strain is at the microcolony level and we use the fatty acid intensity normalized to protein intensity to represent microcolony level production.

Segmentation and tracking of phase contrast images was performed using the DeLTA 2.0 pipeline (62). Segmentation errors were corrected manually prior to downstream analysis. We calculated growth rate of single cells using the logarithmic derivative of cell length with the following formula:

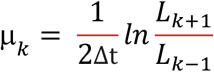

Where μ is growth rate, k is the current frame, Δt is the time between frames, and L is cell length.

### Phase contrast imaging

Cells were imaged with a Nikon Ti-E microscope using a 100x objective with phase contrast imaging. Images were collected every 20 minutes with the microscopy chamber held at 31°C. Production strains were grown on agarose pads containing M9 minimal media as described above for SRS imaging. After 18 hours of growth, the position of the tracked microcolony was recorded and the slide was moved to the SRS microscope for endpoint hyperspectral imaging.

### Stress responsive reporter strain

Cells containing the stress reporter plasmid pBbE-ibpAB-mRFP1 were grown on agarose pads. The cells were allowed to recover on the agarose pads for 3 hours at 31°C prior to SRS exposure. After recovery, a field of view on the pad containing several microcolonies was subject to SRS scanning at various step sizes (150 nm or 230 nm) with power held at 25mW for the Stokes laser and 15mW for the pump laser. Red fluorescent protein (RFP) images were taken of the scanned field of view and a nearby, un-scanned field of view every 30 minutes. Since the RFP is photobleached from the SRS scan, the change in RFP of each microcolony was calculated for each condition. To account for focus differences between fluorescent images at different time points, the scanned field of view was normalized to the RFP of the nearby, un-scanned microcolonies.

## Acknowledgements

This work was supported by the Office of Science (BER) at the U.S. Department of Energy (DE-SC0019387 to MJD, JXC, WWW), the National Science Foundation (1804096 to MJD), and NIH R35 GM136223 to JXC. We thank Dr. Normal Lee and the Chemical Instrumentation Center for assistance with GC-MS experiments. Dr. Joshua Finkelstein provided valuable input on the manuscript.

## Author Contributions

N.T. and H.L. performed experiments and conducted data analysis. M.J.D., J.X.C., and W.W.W. provided overall guidance on the project. N.T. was responsible for strain construction, production quantification, and sample preparation. H.L. performed SRS imaging. J.B.L. performed pilot experiments with a production strain. O.M.O. helped with single-cell segmentation and tracking of phase contrast imaging experiments. D.B. helped to develop the GC-MS protocol and quantified strain production. N.T., H.L., and M.J.D. wrote the manuscript with input from J.B.L., W.W.W., and J.X.C.

## Data Availability

The datasets generated during and/or analyzed during the current study are available from the corresponding author on reasonable request.

## Code Availability

The code for spectral analyses used in this study is available from the corresponding author on reasonable request.

## SUPPLEMENTARY INFORMATION

### Supplementary Figures

**Figure S1.**
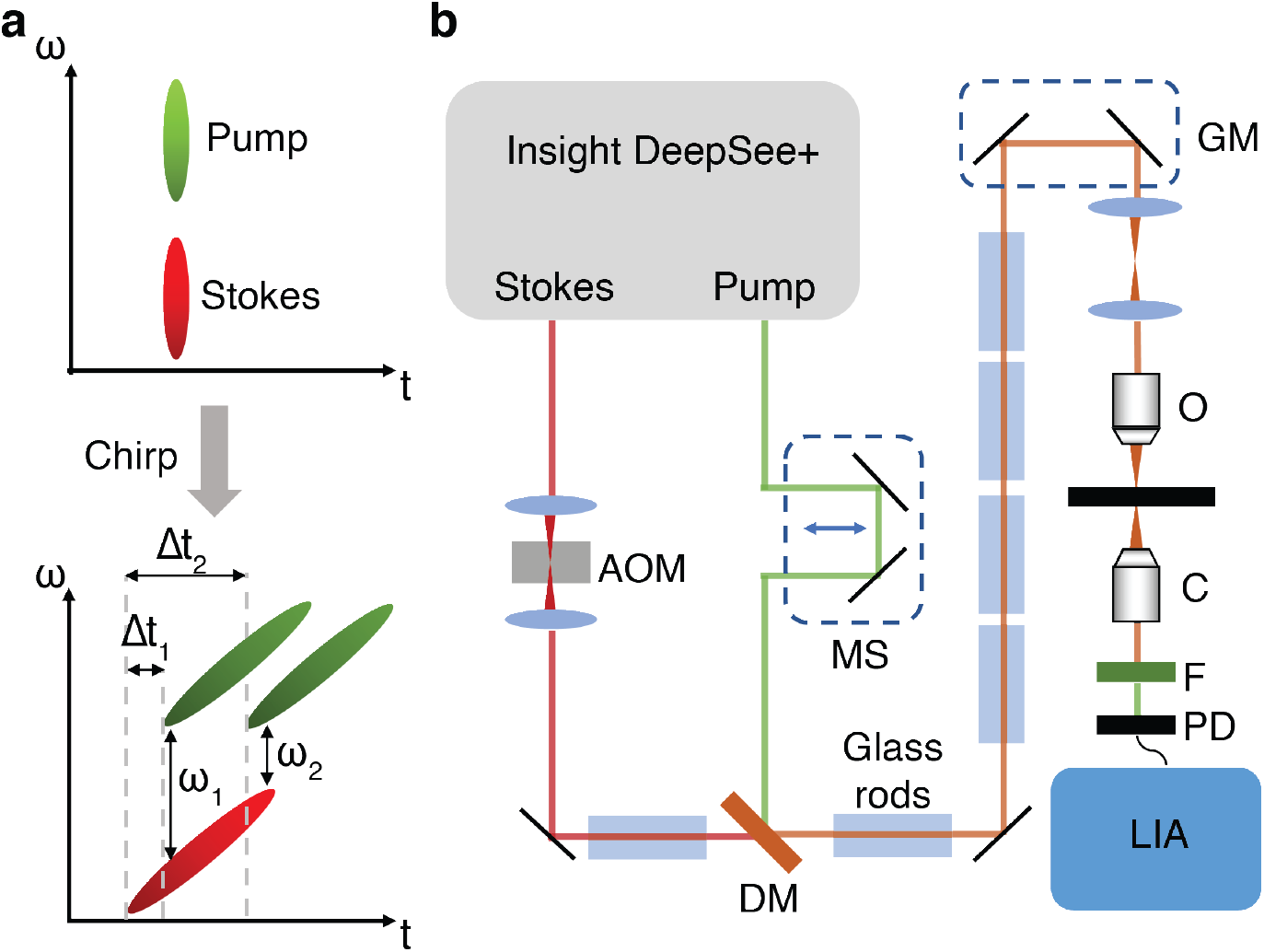
Hyperspectral SRS setup. **(a)** Concept of hyperspectral SRS using spectral focusing. The pump and Stokes lasers are linearly chirped by high dispersion glass rods to temporally separate the spectral components. Each temporal delay between the two pulses corresponds to a Raman vibrational mode. **(b)** Optical setup. AOM, acousto-optic modulator; MS, motorized stage; DM, dichroic mirror; GM, galvo mirrors; O, objective; C, condenser; F, filter; PD, photodiode; LIA, lock-in amplifier.

**Figure S2.**
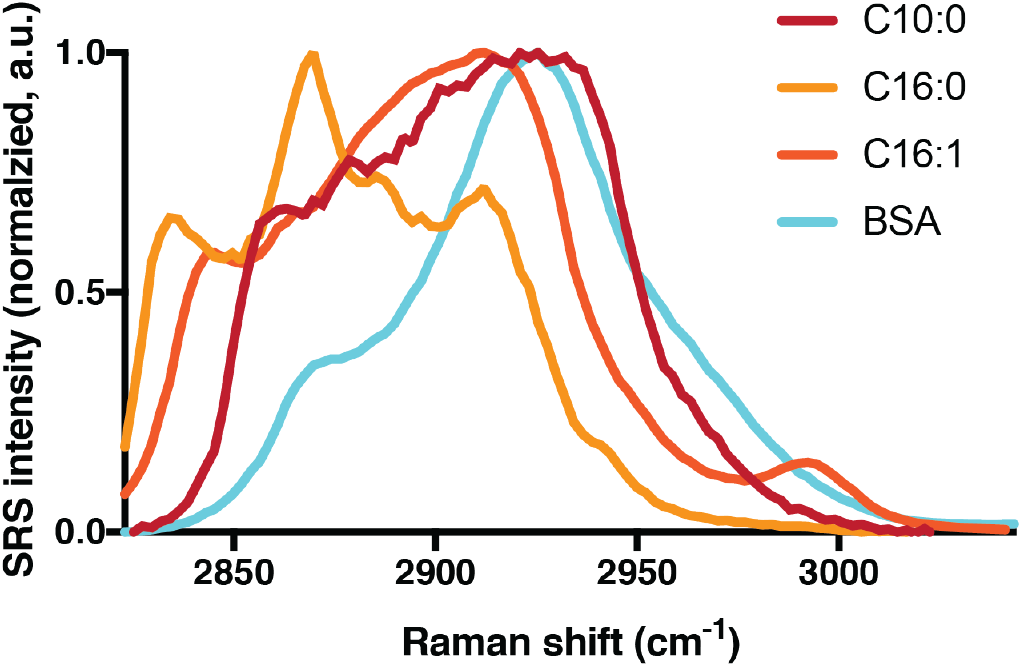
SRS spectra of pure standards used to analyze hyperspectral images to produce chemical maps. (BSA: bovine serum albumin, C10:0: decanoic acid, C16:0: palmitic acid, C16:1: palmitoleic acid).

**Figure S3.**
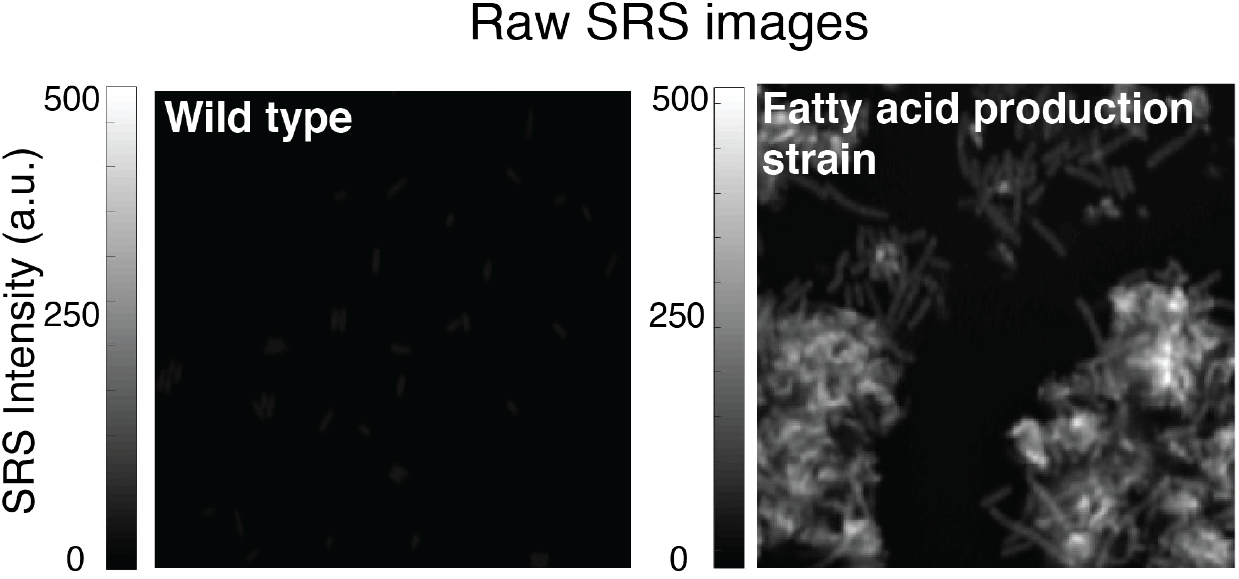
Raw SRS images shown in Fig. 1c of wild type and a strain overexpressing a cytosolic thioesterase (*Ab*TE*), but with both images scaled with the same color axis for direct comparison.

**Figure S4.**
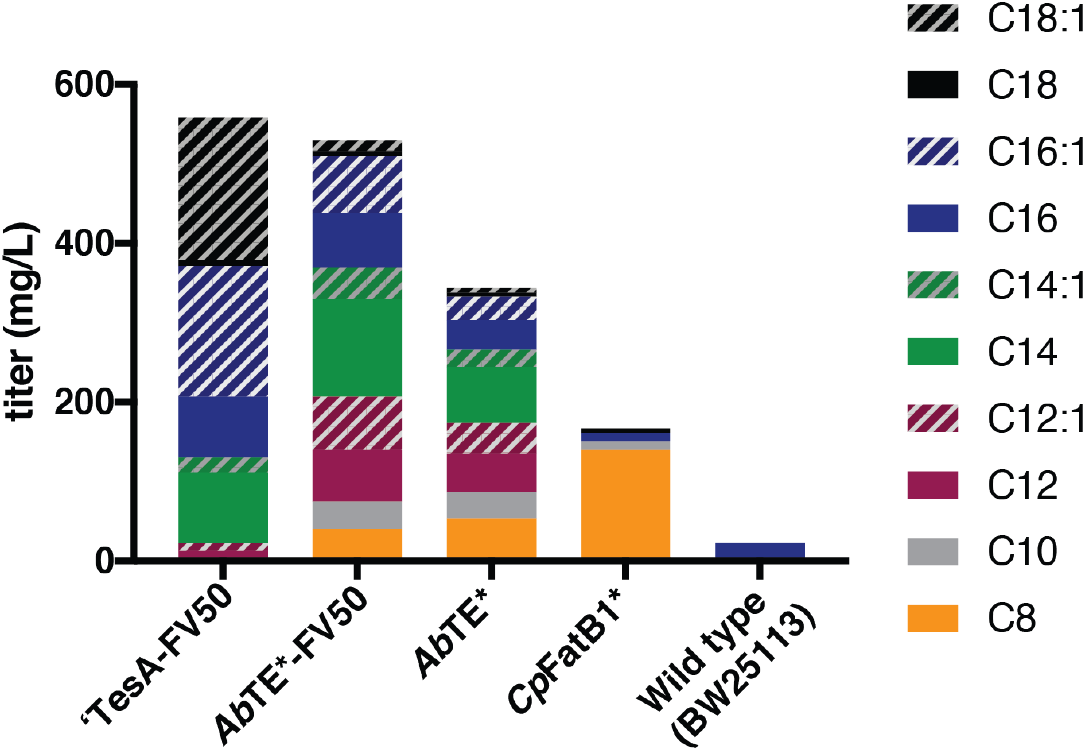
Fatty acid production quantification for strains in this study. GC-MS quantified fatty acid production data for each strain. Cells were grown 24 hours post thioesterase induction in liquid culture. For chain length prediction, these exact cultures were taken for SRS imaging at the same timepoint.

**Figure S5.**
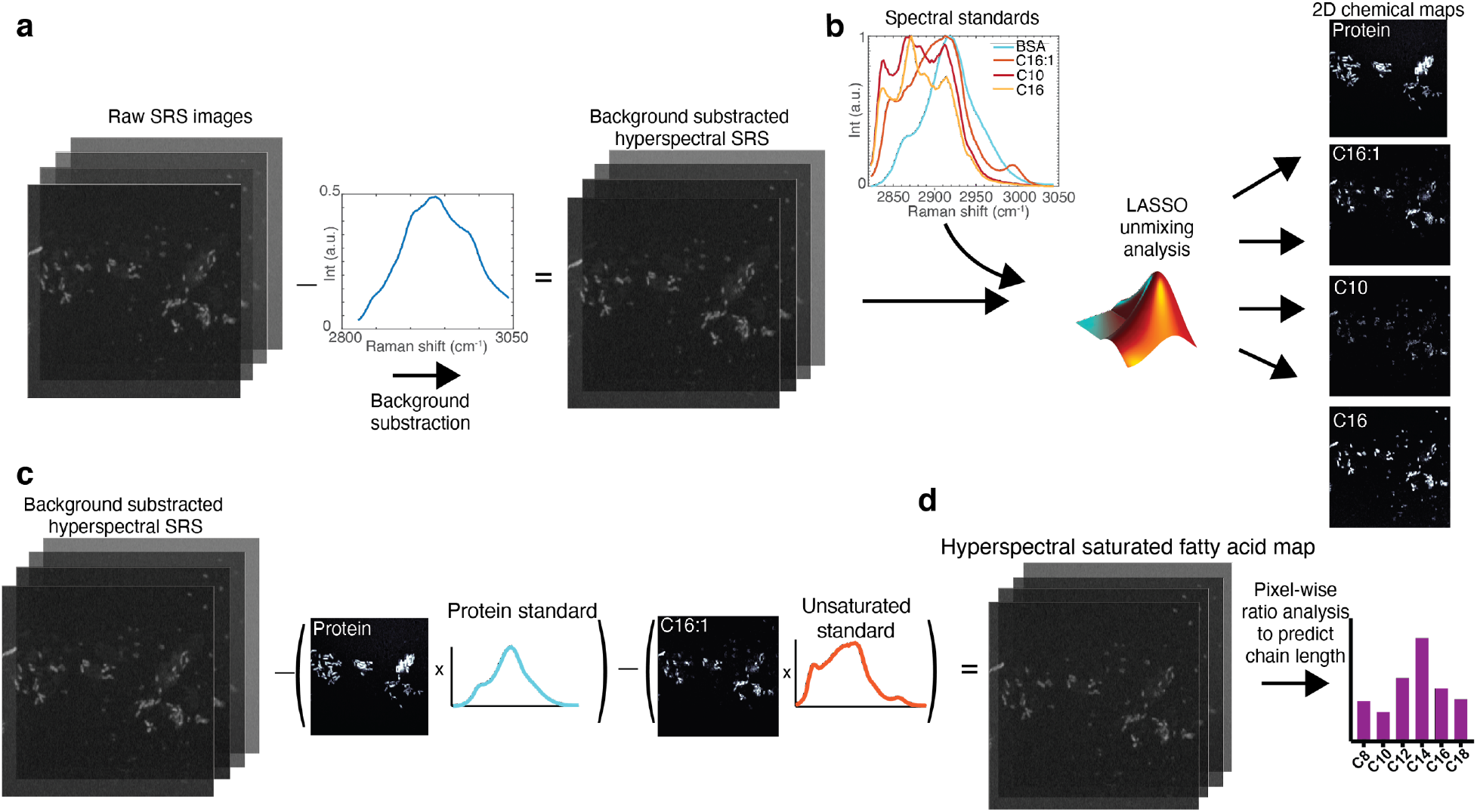
Analysis workflow for chain length prediction from hyperspectral SRS images. **(a)** Raw SRS images are first background subtracted. **(b)** Background subtracted images are unmixed using chemical standards. Protein, BSA; unsaturated fatty acid, C16:1; medium chain fatty acid, C10; and long chain fatty acid, C16. C10 and C16 maps are used to represent a mixture of saturated fatty acids. **(c)** Protein and unsaturated fatty acid maps are multiplied by their respective standard spectra and subtracted from the background-subtracted hyperspectral image to produce a three-dimensional saturated fatty acid map. **(d)** Ratio analysis is performed on each pixel to calculate chain length and weighted by raw intensity to predict chain length distribution of the field of view.

**Figure S6.**
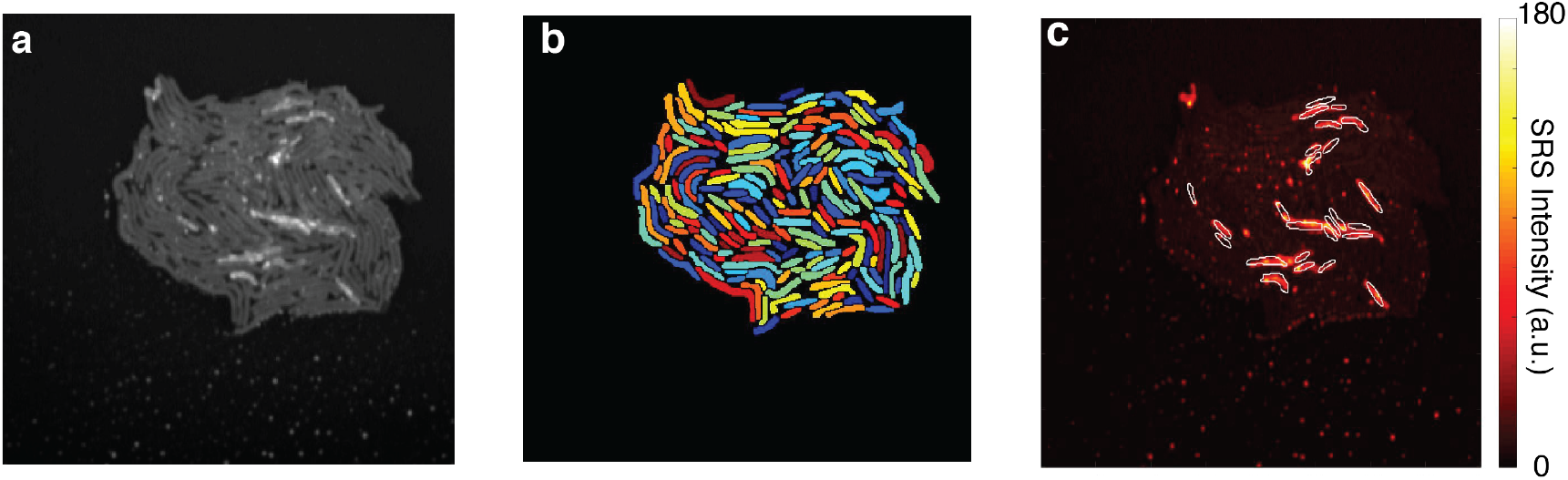
Single cell segmentation of a microcolony. **(a)** Raw SRS images are used to segment microcolonies to perform single cell analysis shown in Fig. 3c. **(b)** Segmentation of microcolony in (a). **(c)** Segmentation of the top 25 highest producing cells overlaid on the fatty acid map of the microcolony.

**Figure S7.**
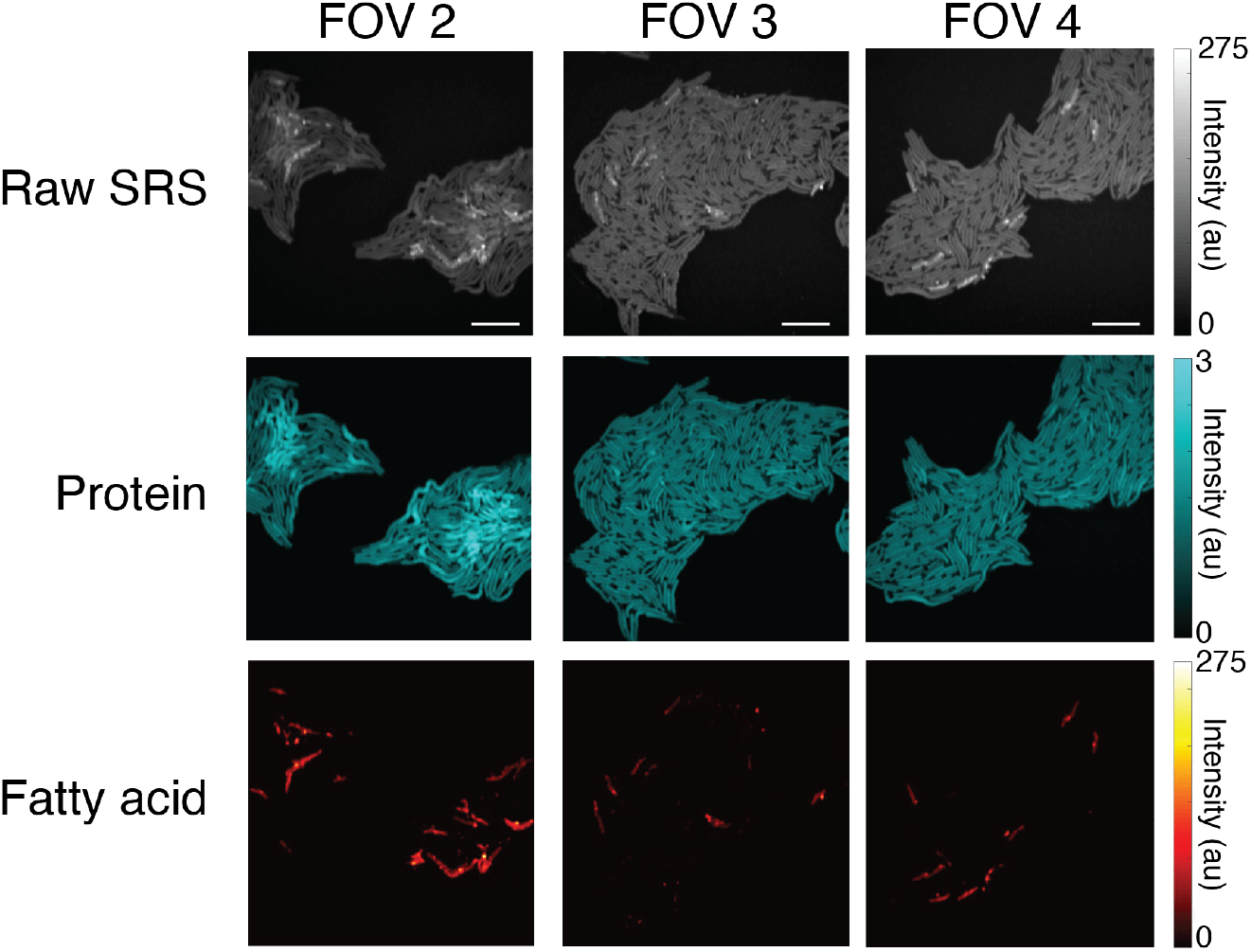
Intra-colony heterogeneity of the *Ab*TE* strain. **(a)** Three additional fields of view (FOV) of the *Ab*TE*-FV50 strain shown in Fig. 3b. Raw SRS, protein, and fatty acid chemical maps are shown for all. Scale bars, 10 μm.

**Figure S8.**
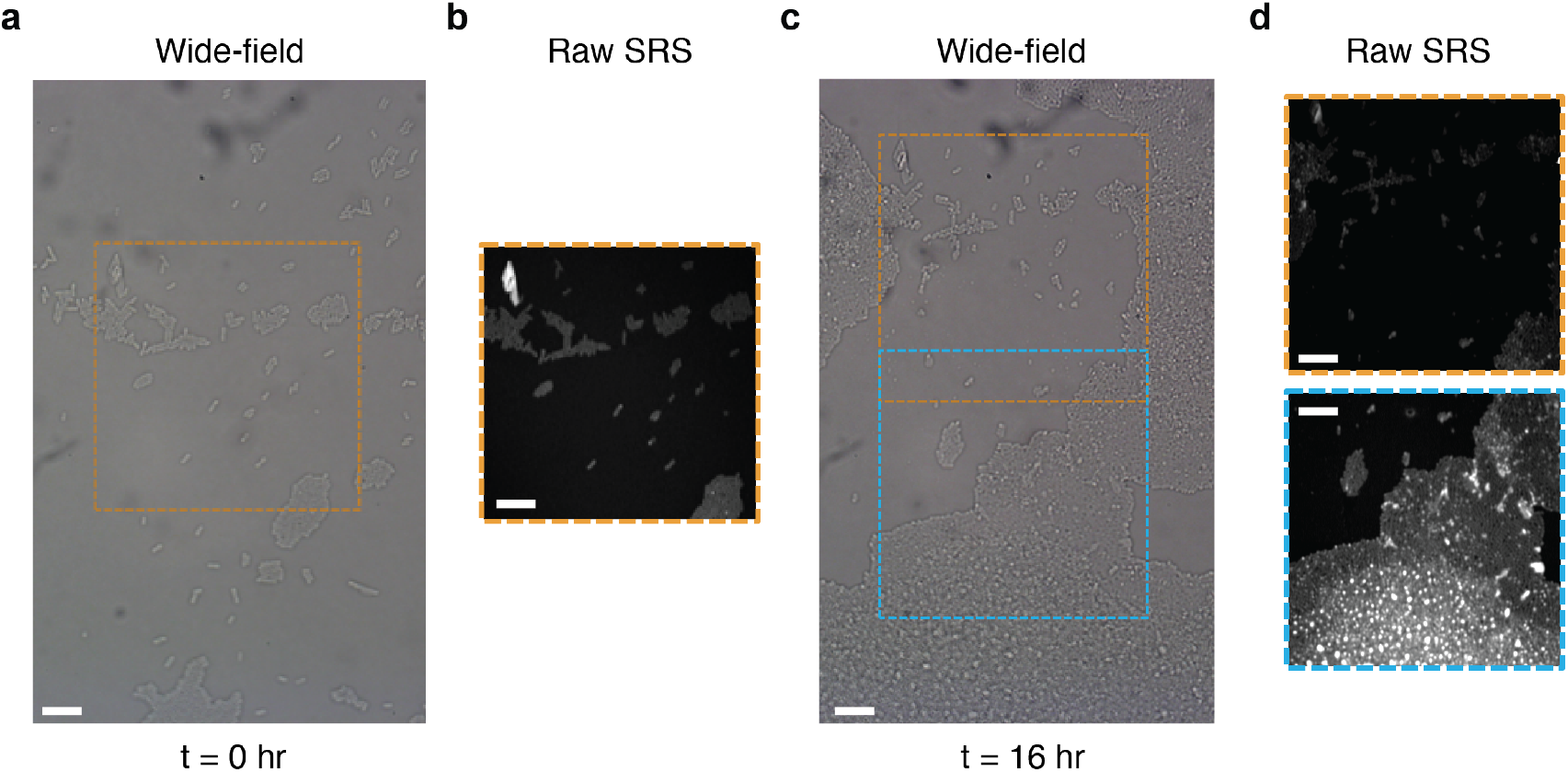
Testing photodamage of live *E. coli* cells. **(a)** Wide-field transmission image of *E. coli* cells at the start of the cell incubation (t = 0 hr). **(b)** Hyperspectral SRS image of the region highlighted with a yellow rectangle in (a). **(c)** Wide-field transmission image of the same field of view after incubation (t = 16 hr). **(d)** Hyperspectral SRS images of the previously scanned region (yellow rectangle in (c)) and an adjacent region without previous SRS laser exposure (blue rectangle in (c)). Scale bars, 10 μm.

**Figure S9.**
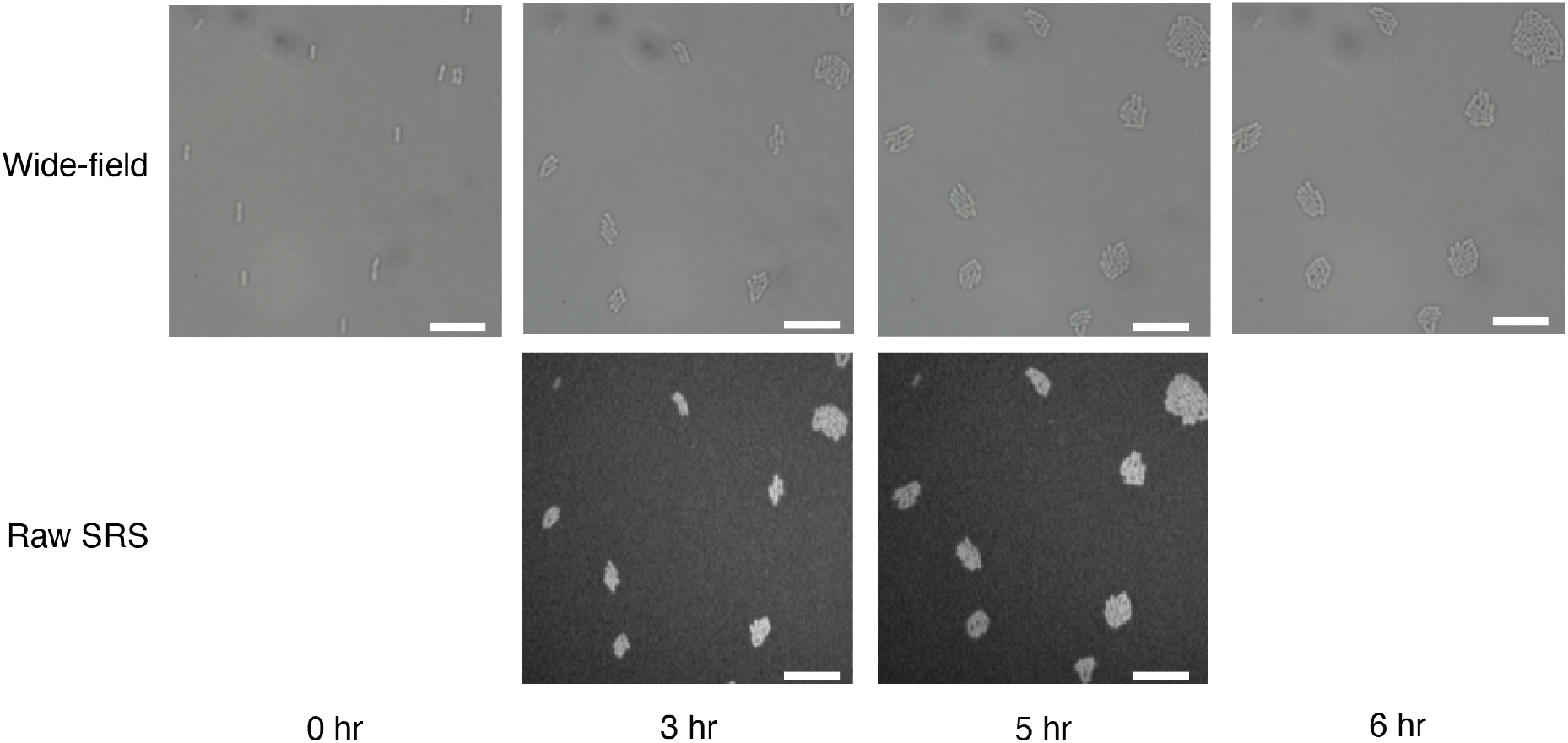
Optimized SRS laser powers enable live cell imaging of *E. coli*. Wide-field transmission image of *E. coli*, with raw hyperspectral SRS images of the same region for the t = 3 and 5 hr timepoints. Spectral summation is shown. Scale bars, 10 μm.

**Figure S10.**
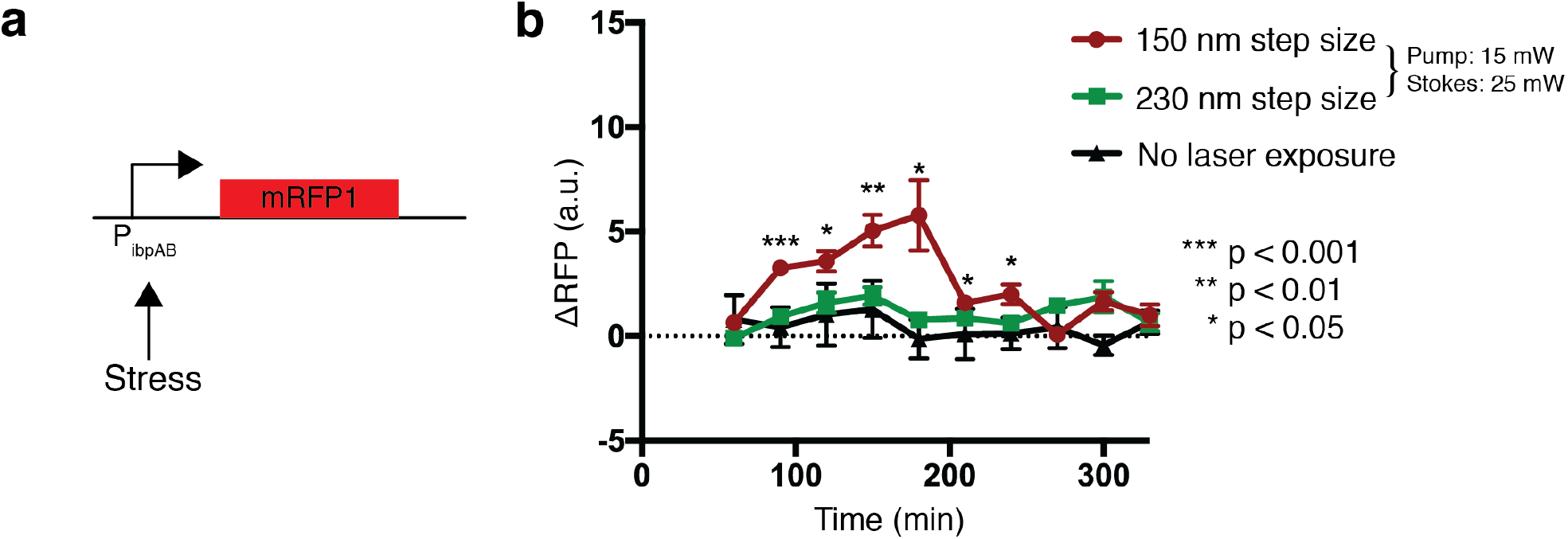
Stress response of longitudinal SRS imaging conditions. **(a)** Schematic of stress reporter, P_ibpAB_, driving expression of mRFP1. **(b)** Fluorescent response of cells containing the reporter after SRS exposure. Low power SRS (15mW pump and 25 mW Stokes) was tested using steps sizes of 150nm and 230nm. P-values compare 150nm step size to no laser exposure (n = 9; two tailed unpaired t-test). Error bars show standard error of the mean.

**Figure S11.**
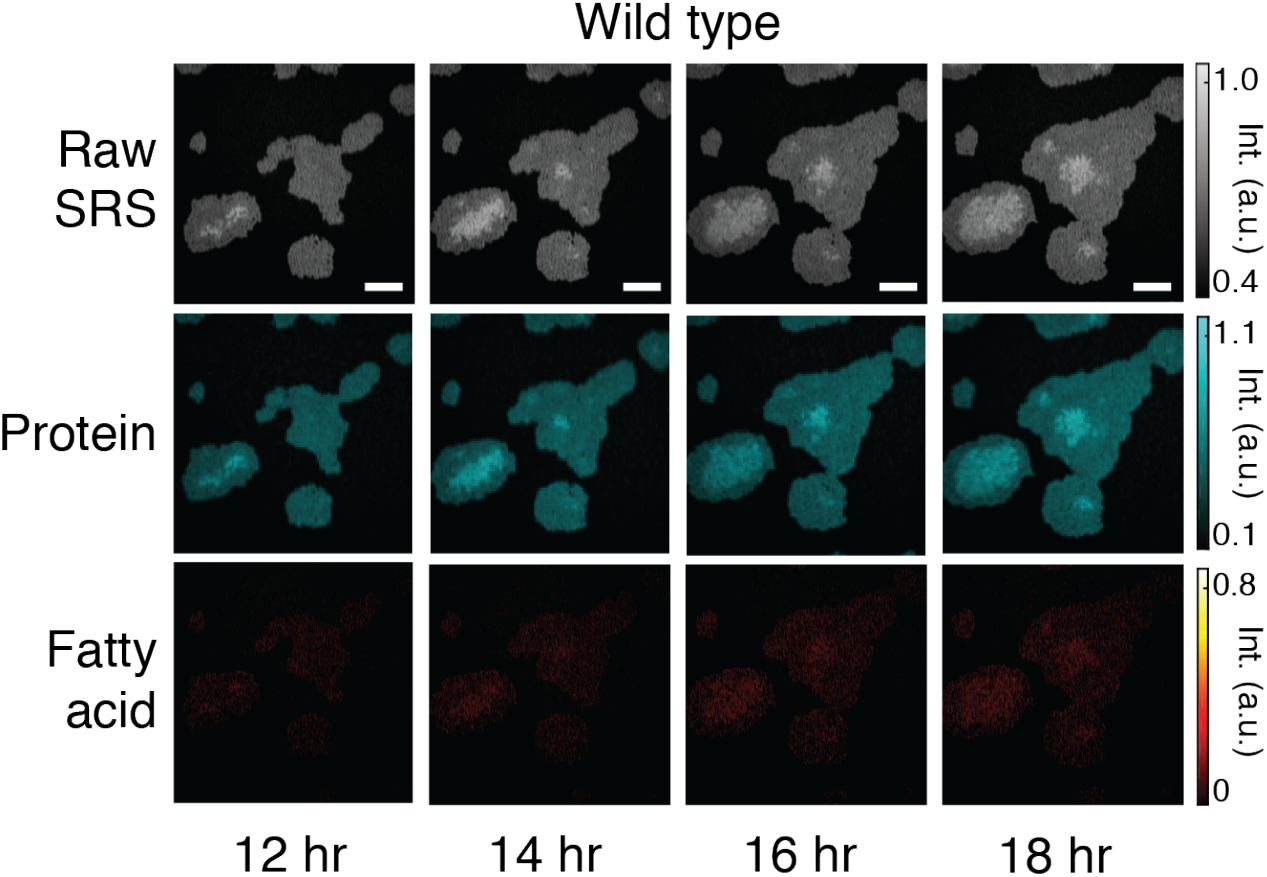
Time-lapse images of a wild type control strain, shown with the raw SRS images (spectral summation of the SRS image stack) and chemical maps corresponding to protein and fatty acid content.

**Figure S12.**
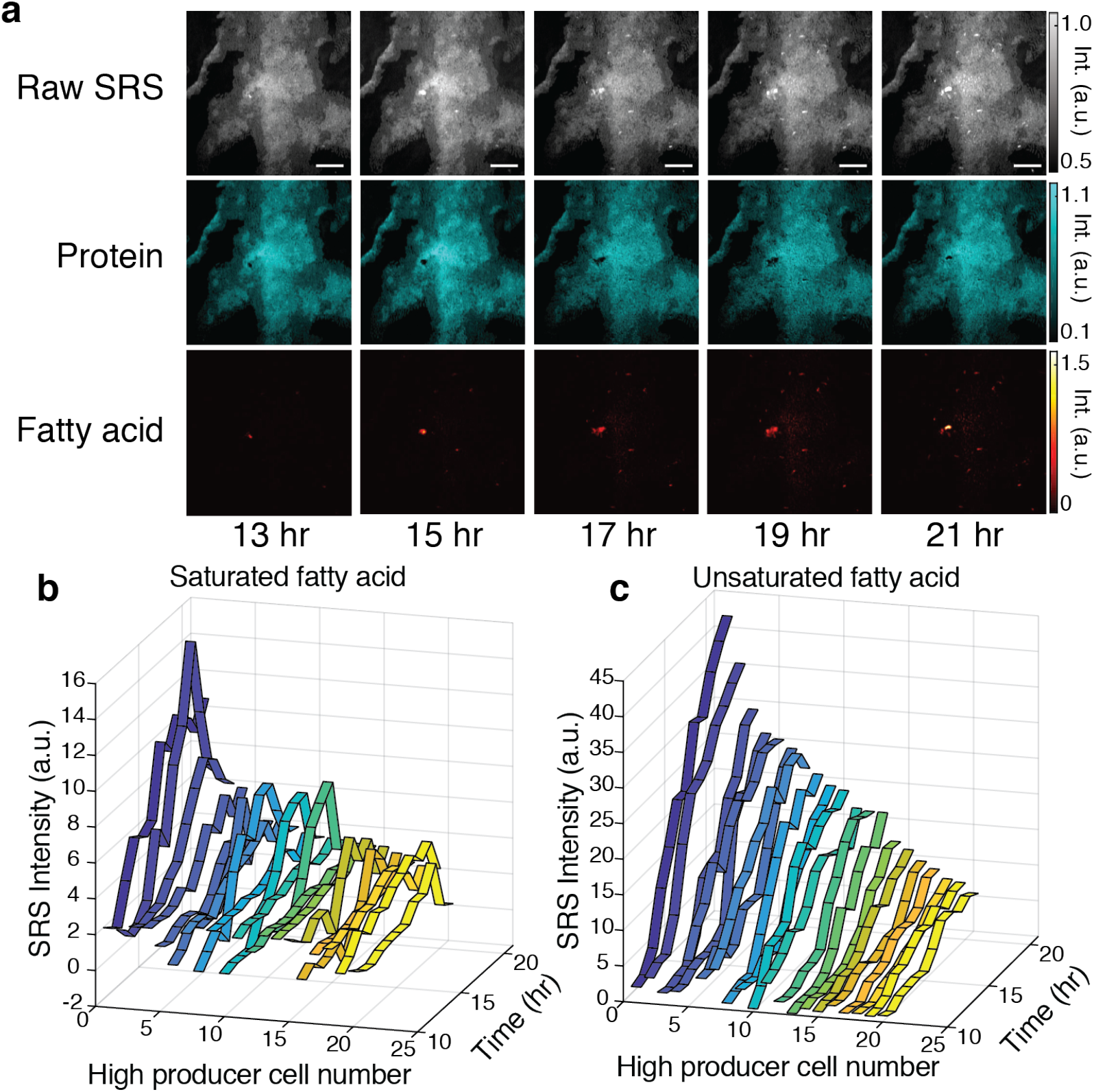
Time-lapse images of fatty acid production in the *Ab*TE*-FV50 strain. **(a)** Raw SRS images, protein, and fatty acid chemical maps are shown. Time values represent time grown on the agarose pad after IPTG induction. Scale bars, 10 μm. (b) Saturated and (c) unsaturated content of high producer single cells from the time-lapse images in (a).

**Figure S13.**
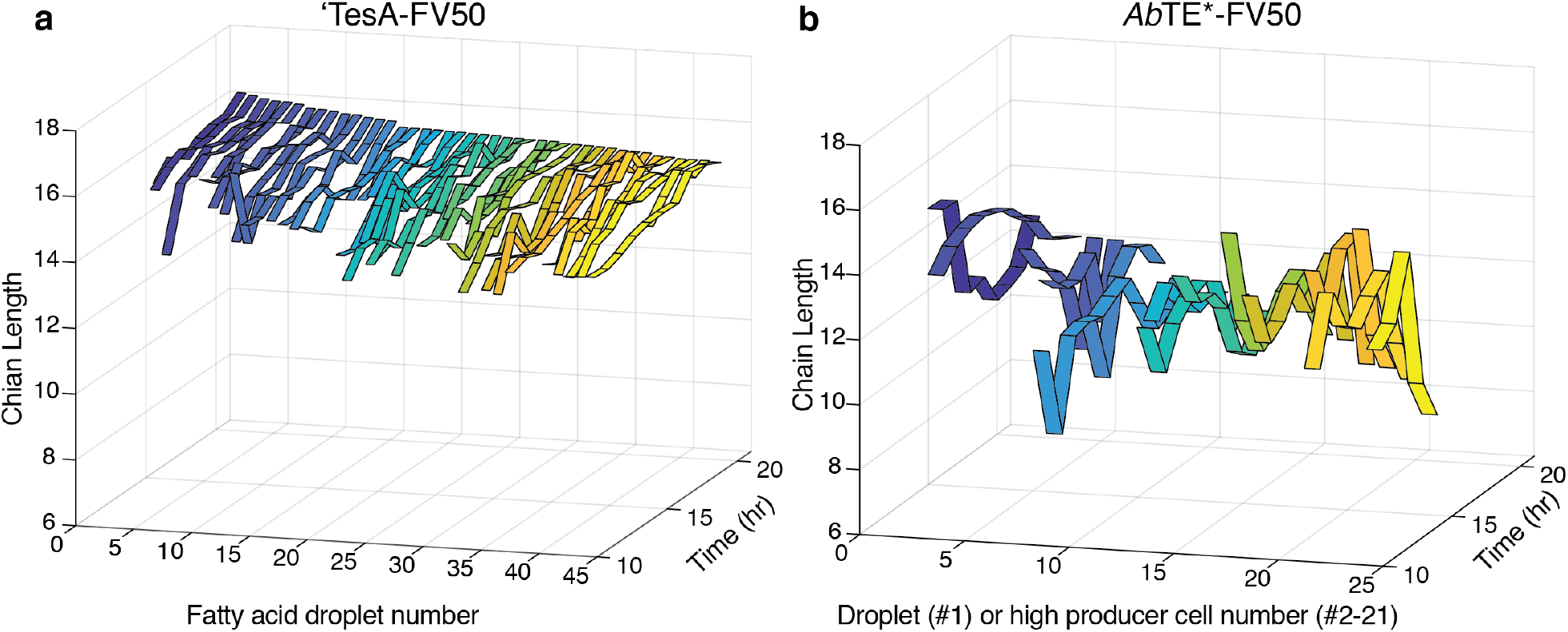
**(a)** Longitudinal chain length predictions of droplets from the ‘TesA-FV50 high microcolony from Fig. 4a. **(b)** Longitudinal chain length predictions of the large droplet (ribbon #1) and high producing cells (ribbons #2-21) in the *Ab*TE*-FV50 microcolony from Fig. S12.

**Figure S14.**
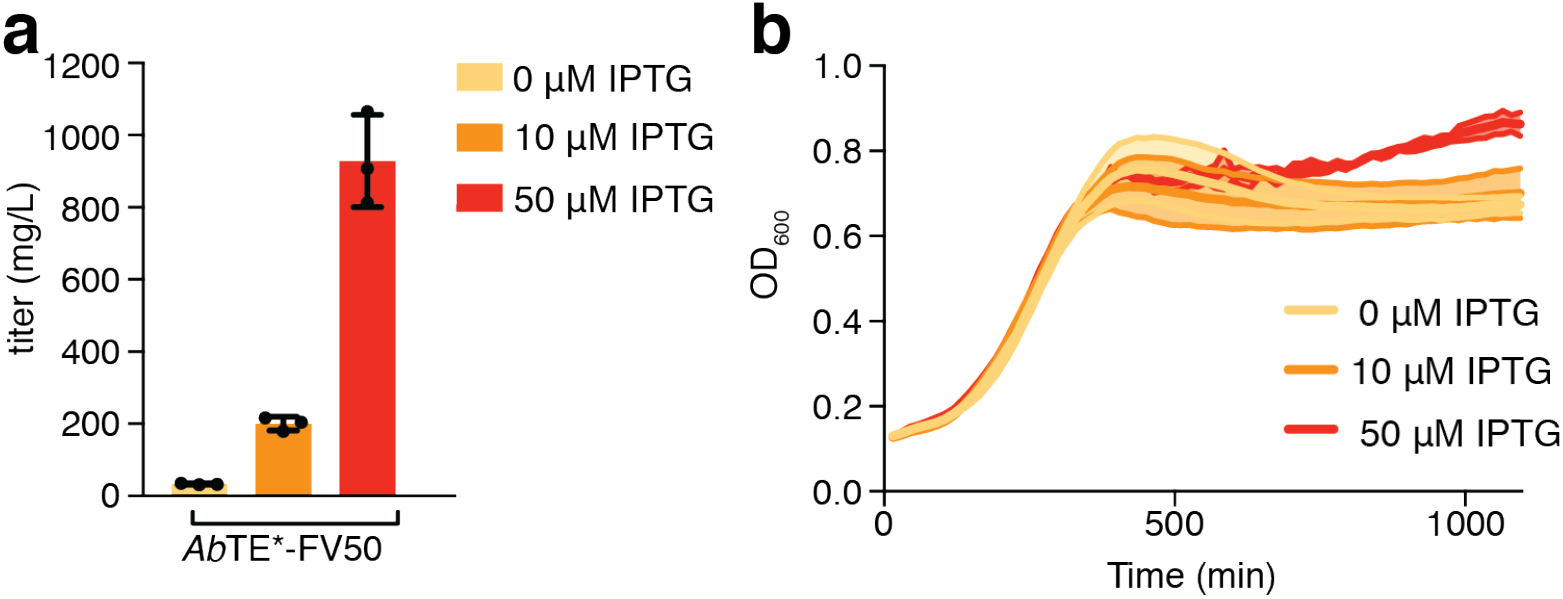
**(a)** GC-MS quantification of fatty acid production and **(b)** growth of *Ab*TE*-FV50 at varying IPTG induction levels (n = 3). Error bars, standard deviation.

**Figure S15.**
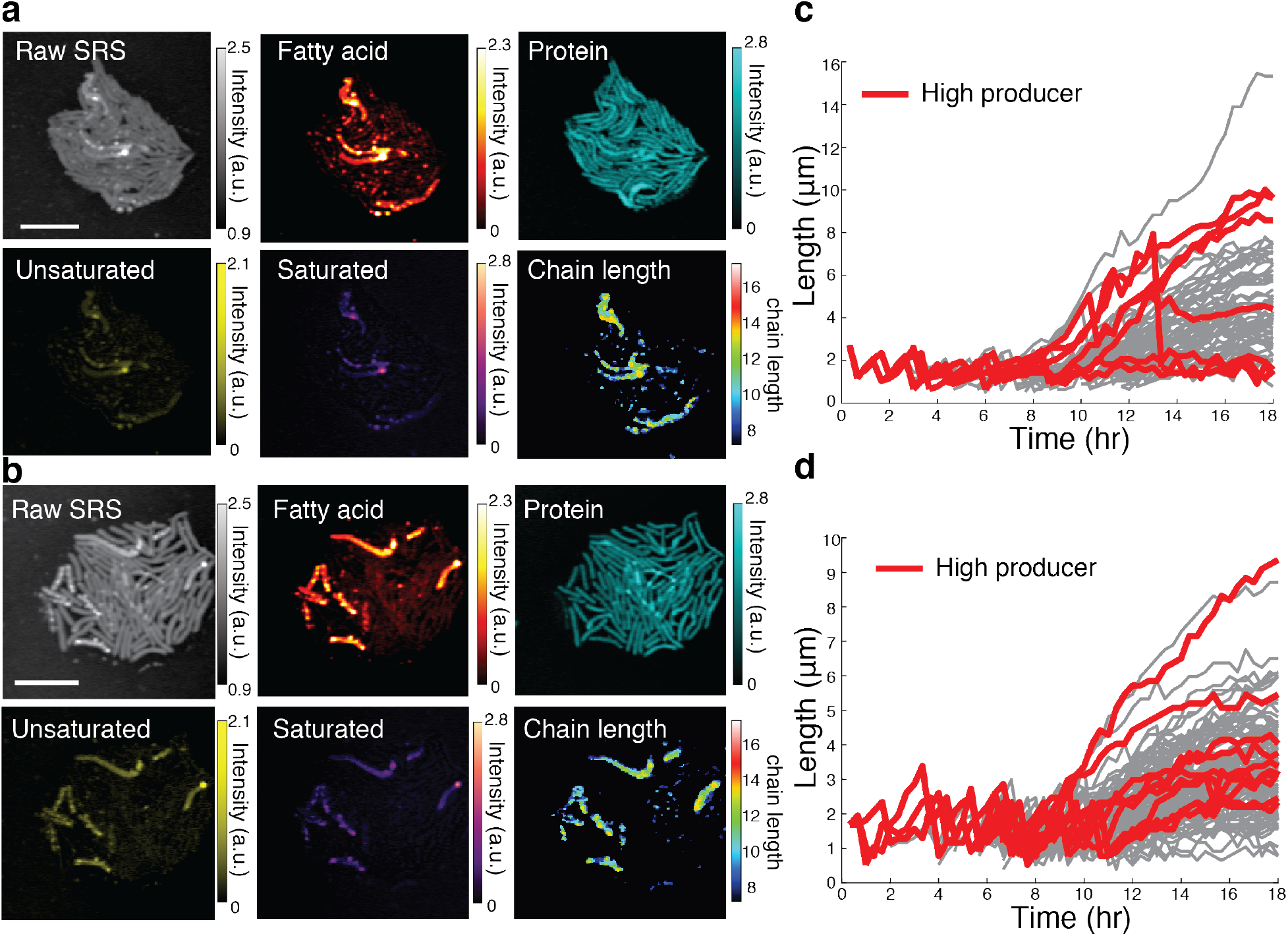
**(a-b)** Endpoint SRS imaging and spectral decomposition of *Ab*TE*-FV50 microcolonies tracked with time-lapse phase contrast imaging. **(c-d)** Single-cell lengths of individual cells in (a-b), with high producer trajectories (top 15%) highlighted in red.

**Figure S16.**
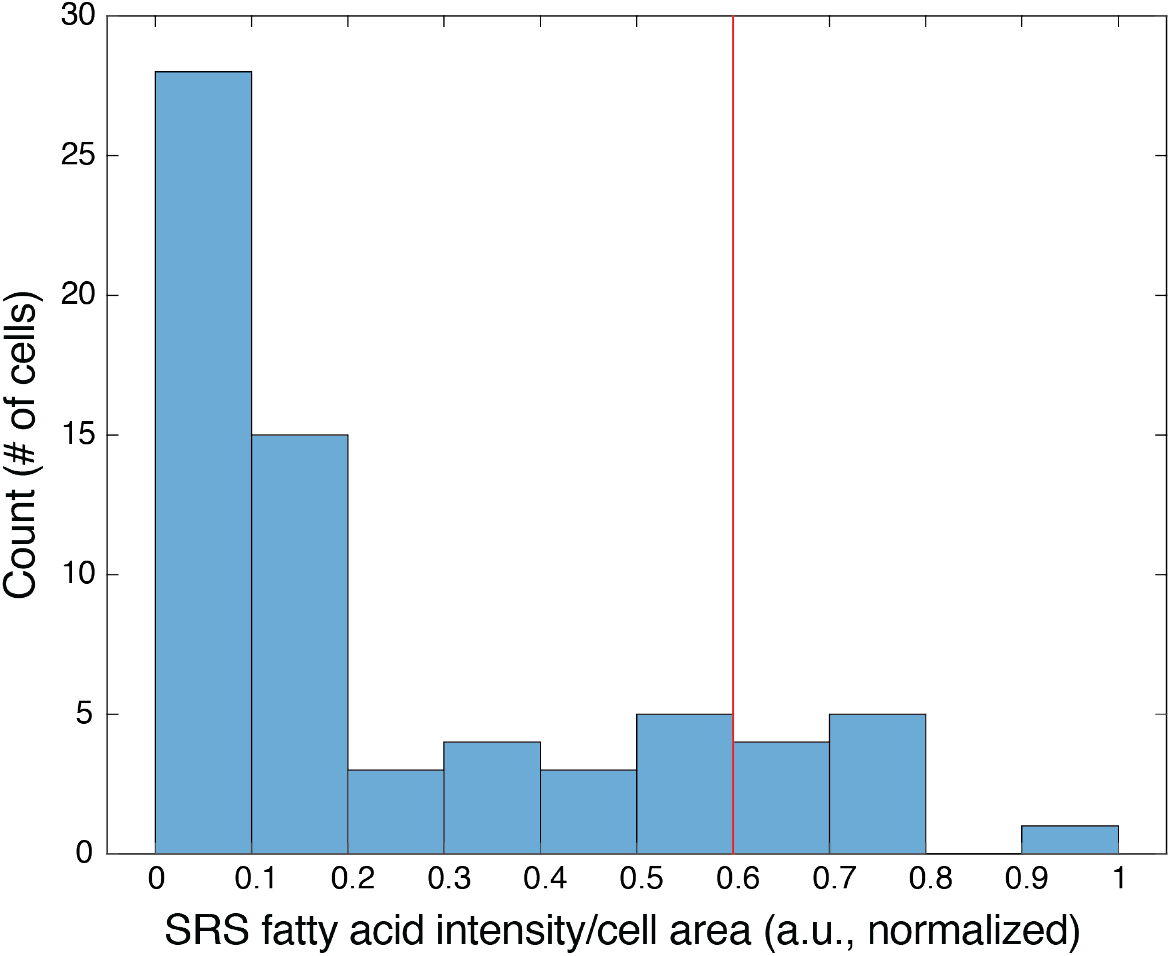
Endpoint fatty acid distribution of the *Ab*TE*-FV50 microcolony in Fig. 5. The red line indicates the threshold set to define high producer cells.

### Supplementary Movies

**Movie S1**. Time-lapse wide-field transmission images of the wild type strain during the live cell SRS imaging shown in Fig. 3c. The white box indicates the SRS imaging region.

**Movie S2**. Time-lapse phase contrast images of the *Ab*TE*-FV50 microcolony from Fig. 5.

**Movie S3**. Time-lapse phase contrast images of the *Ab*TE*-FV50 microcolony from Fig. S15a.

**Movie S4**. Time-lapse phase contrast images of the *Ab*TE*-FV50 microcolony from Fig. S15b.

**Movie S5**. Manually segmented droplets of the ‘TesA-FV50 strain used for compositional tracking in Fig. 4e-f and Fig. S13a.

**Movie S6**. Manually segmented droplets of the *Ab*TE*-FV50 strain used for compositional tracking in Fig. S12b-c and Fig. S13b.

